# Correlated Gene Copy Number Changes in a Seminal Fluid Protein Network in *Drosophila*

**DOI:** 10.1101/2025.10.27.684572

**Authors:** J.A. Carlisle, B.K. McCormick, L. Verbakel, A.C. von Philipsborn, M.F. Wolfner, A.G. Clark

## Abstract

Reproductive proteins often diverge rapidly between species, yet network function must be maintained. Shared selective pressures on network members and compensatory changes between members can drive their parallel evolutionary trajectories. Indeed, correlated evolutionary rates of amino acid sequence change have been observed for interacting reproductive proteins. But whether gene copy number changes also correlate has not been widely studied. Here, we investigated copy number variation (CNV) of genes in the *Drosophila* Sex Peptide Seminal Fluid Protein (Sfp) network. Previous research analyzed CNV of the Sfp Sex Peptide (SP) in *Drosophila* species. We focus on 9 other Sfps whose function is required to mediate the binding of SP to sperm in *D. melanogaster* which is required for persistence of female post-mating responses. To exhaustively annotate CNV of genes, we developed a computational pipeline pairing iterative protein queries to genome sequence searches with phylogenetic clustering to resolve homology relationships. We observed that the Sfp network’s genes are ancestral to *Drosophila* and that there were repeated duplications and losses of network members across the genus. We detect statistically significant correlations in gene duplication or loss events among network proteins, and show this can be used to identify new members of the network. We also investigated CNV of female-derived proteins that act downstream of the SP sperm-binding network to modulate SP function, these proteins showed no significant correlation of gene turnover events with SP or its network. Our results provide insight into how evolving reproductive genes tolerate duplication and loss, and how network relationships could constrain reproductive protein evolution.

**Significance Statement:** Reproductive proteins often diverge rapidly between species, yet network function must be maintained. Shared selective pressures on network members and compensatory changes between members can drive their parallel evolutionary trajectories. We report correlated gene duplication and loss among members of the *Drosophila* Sex Peptide seminal fluid protein network, suggesting that duplication or loss events may drive corresponding events in other network genes. This work is a natural extension of the idea of evolutionary rate covariation, but instead of scoring rates of substitution it tracks correlated duplication and loss events on the phylogeny. Applied to the Sex Peptide network, the method reveals striking patterns, especially for coordinated loss, and identifies a new network gene that is experimentally confirmed.

## Introduction

Reproductive proteins frequently show remarkable divergence between even closely related species, a phenomenon observed from mammals to invertebrates to plants and to sexually reproducing microbes^1–5^. Sexual selection and sexual conflict are frequently cited as explanations, although in some cases rapid change may instead reflect relaxed selective constraints ^1,6–9^. Shared selective pressures on members of gene networks, paired with compensatory changes of network members to preserve function, can drive parallel evolutionary trajectories of network members ^10–13^. Indeed, this coordinated evolution of functionally related genes has been documented for gene expression and sequence variation ^10,14^. Yet, whether such coordination extends to gene copy number is largely unknown. We hypothesize that co-divergence in copy number among functionally related genes can provide insight into how genes operate together to mediate reproduction. Correlated patterns of gene copy number could even be used to identify new network members, in a guilt by association approach. Here, we test this hypothesis by investigating patterns of coordinated gene copy number variation (CNV) of members of a seminal fluid protein network that controls the long-term post-mating response in *Drosophila melanogaster*.

Seminal fluid proteins (Sfps) are key mediators of reproductive success, influencing sperm storage, fertilization, and female post-mating physiology ^15,16^. *D. melanogaster* is a premier genetic model for studying these functions, and the evolution of *Drosophila* Sfps is often cited as a prime example illustrating the evolution and rapid diversification of reproductive genes ^8,17–20^. As phylogenetic distance increases, the number of recognizable *D. melanogaster* Sfps declines sharply, with only ∼15% detectable in the accessory gland proteome *D. virilis* ^21,22^. This loss of orthologs likely reflects a combination of rapid sequence divergence, gene turnover, and paralogy that obscures orthology. The recent availability of high-quality genome assemblies across the genus *Drosophila*, along with improved phylogenetic methods, provides an opportunity to improve the resolution of homologous relationships across evolutionary history and map the evolutionary trajectories of copy number changes of Sfps ^23–28^. On average, Sfps show higher sequence divergence than most other non-reproductive proteins ^8,18^. Beyond the evolutionary pressures that promote diversification of reproductive proteins, Sfps may also evolve more rapidly than intracellular proteins because, as secreted proteins, they experience fewer structural constraints and are exposed to more dynamic extracellular interactions ^29–31^. Further, many Sfps also show expression restricted to the reproductive tract, reducing pleiotropic constraint on their evolution ^31^. These factors may explain why, for some Sfps, copy number has also been observed to vary dramatically between species, making Sfps an exceptional system for probing the evolutionary forces driving CNV in reproductive genes and for testing how covariation in copy number may manifest within reproductive networks.

In *D. melanogaster*, the Sfp Sex Peptide (SP) regulates varied and persistent post-mating changes in female physiology and behavior, both reproductive and non-reproductive, through interaction with its female receptor SPR (Sex Peptide Receptor) ^32–34^. These effects persist long-term and requires SP binding to sperm ^35^. A network of Sfps is necessary for this sperm binding and thus for the female long-term post-mating response (LTR) ^13,34,35^. The LTR includes the female exhibiting a sustained reduction in receptivity and increased egg laying ^35^. The Sex Peptide Network (SPN) consists of Sfps that act after sperm enter the female and before they reach the sperm storage organs to enable stabilization of SP bound to sperm and localization of sperm to storage organs (see also Figure 1) ^35^. Known SPN members belong to protein families (serine proteases and homologs, CRISPs/CAP-domain proteins, lectins) that are also found in the seminal fluid of species across the tree of life, including humans ^15,36–43^. Successful storage of sperm with bound SP, followed by gradual release of SP’s active region from sperm, is essential to sustain the female’s long-term post-mating response ^35^. Additional female-derived factors further modulate SP function and SP’s binding to sperm; however, the identities of the relevant female reproductive tract molecules remain unknown ^44^.

**Figure 1.**
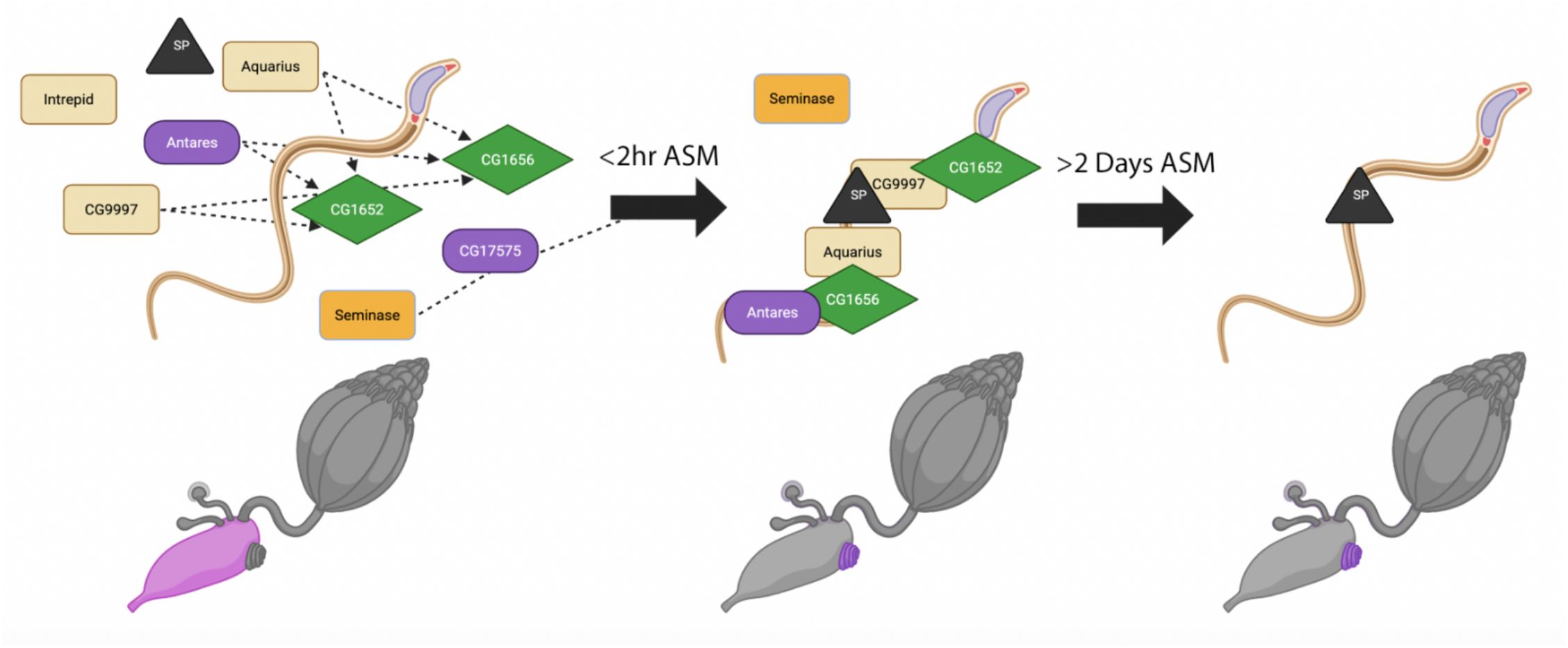
Seminal fluid proteins in the Sex Peptide Network are critical for Sex Peptide binding to sperm and for SP’s long-term effects on female reproduction. Figure adapted from Allen 2024^97^. While all members of the Sex Peptide Network (SPN) are critical for storage of sperm with bound Sex Peptide (SP), there is evidence that these proteins mediate different steps of a cascade of events that allow for the long-term effects of SP^35^. These long-term effects are mediated through SP’s long-term binding to sperm stored in the seminal receptacle, the primary sperm storage organ. Some of these seminal fluid proteins (Sfps) (CG1652 and CG1656) require other network members (Aqrs, Antr, CG9997) for proper transfer to the female. However, while essential for transfer of CG1652 and CG1656, Antr’s transfer is independent of other known network members. After being transferred to the female, Sems and CG17575 do not bind sperm nor are stored in the seminal receptacle, but they are critical for SP and CG1656 to localize to the seminal receptacle where both proteins (along with CG1652, Antr, and CG9997) bind sperm. Sems can be detected in the seminal receptacle despite not binding sperm. While SP binds to sperm long-term, its network does not. Most of the SPN Sfps are no longer detectable on sperm 1 day after mating and none are detectable after 4 days although SP can still be found bound to sperm^35,98^. The functions of all SPN Sfps have not been fully characterized. For example, Intr’s function within the network has not been described, and additional members of the network had yet to be identified. Two of the SPN Sfps, *CG1652* and *CG9997*, evolved under positive selection^46^. Of the 8 known members of the SPN, 4 are serine proteases, 2 are lectins, and 2 are proteins containing a CAP domain. Seminal fluid proteins across species, including in mammals, have been shown to contain these protein domains. Known interactions are indicated by dashed lines. The first panel depicts transfer of Sfps and sperm during copulation into the bursa, the second panel depicts SP and SPN proteins bound to sperm 2 hours after the start of mating, the third panel depicts only SP detectable as bound to sperm more than 2 days after mating. Orange squares are serine protease homologs of the SPN with the darker square being an active protease, purple ovals are CAP domain-containing proteins, green diamonds are lectins, and the black triangle is SP. Figure made with BioRender.

SP originated within *Drosophillidae* in the ancestor of the subfamily *Drosophilinae*, which includes the genus *Drosophila* ^12,32^. SP has no known homologs outside *Drosophilinae* subfamily, however its C-terminus, which is responsible for stimulating the LTR in *D. melanogaster,* is highly conserved^12,45,46^. SP shows extensive variation in copy number across the genus *Drosophila*, with some species having up to 7 copies^45^. In particular, higher copy number is observed within the clade where SPR acquired expression in the female reproductive tract^47^. Given that in *D. melanogaster* the SPN is critical to SP’s long-term function in the female LTR, it may be expected that network Sfps may share similar selective pressures across the genus *Drosophila*, inducing signatures of coevolution between them^32,35^. Indeed, SP shows correlated evolutionary rates in its protein sequence with some, though not all, members of its network^13^. Such correlated patterns of sequence evolution have been used to identify several SPN members through Evolutionary Rate Covariation (ERC) with SP or with other previously identified network proteins^13^. Therefore, it seemed that the SPN would be an ideal testing ground to investigate correlations in copy number changes between functionally related reproductive proteins. We traced the origin of members of the SPN and their CNV across the genus Drosophila. We detected correlated duplication events or “gains” of SP with one member of its network. Interestingly, while loss events were strongly correlated between SPN proteins, no clear association was observed between SP loss and loss of network members.

## Materials and Methods

### Construction of phylogenies of *D. melanogaster* protein families

Paralogous genes within the species *D. melanogaster* were identified using Flybase annotations^48,49^. Translated protein sequences were acquired from FlyBase Batch Download using Fbgn identifiers^48^. Protein sequences were aligned with PROMALS3D^50^. The Interpro plug-in on Geneious (2024 versions) was used to annotate protein domains on sequences from the protein alignment^51^. The alignment was trimmed to the protein domain of interest as defined by Interpro, either a lectin domain (IPR001304), serine protease domain (IPR001254) or CAP domain (IPR014044)^52^. Trimmed protein alignments were used to construct a maximum likelihood phylogeny using RAxML-NG^53^ with 1000 non-parametric bootstraps with resampling or until convergence. The best scoring topology of 20 starting trees (10 random and 10 parsimony-based) was used. Homologous protein relationships established through this analysis of *D. melanogaster* protein family members were validated by phylogenetic clustering of homologs collected from other species. Proteins were determined to be Sfps by following Wigby et al. 2020^15^.

#### Annotation of orthologous and paralogous sequences in 135 Drosophilid species to assess copy number variation

Annotation of genes belonging to each gene family of interest and resolution of orthology and paralogy relationships between them was carried out via iterative blast and phylogenetic clustering (Supplementary Figure 1)^54,55^. The initial tblastn query for each gene family was constructed as follows: for single-exon families, we obtained the full-length amino acid sequences of all genes nested within the family of interest present in *D. melanogaster* and we used tblastn to collect the five best hits (sorted by e-value, with cutoff: 1e-5) for each of these genes in the 35 Drosophila species for which NCBI predicted transcriptomes were available (Supplemental File 1, Table 1). For multi-exon families, the initial query consisted of only the amino acid sequences of the longest exon containing a conserved functional domain for each focal *D. melanogaster* gene.

The resultant query files were then used to identify the coordinates of the five best hits for each query gene in 135 Drosophilid genomes (Supplemental File 1, Table 2) using tblastn (sorted by e-value, with cutoff: 1e-5). Using bedtools, we subsequently merged overlapping hits and obtained the corresponding genomic DNA sequences^56^. Open reading frames were predicted using the EMBOSS getorf tool (see Supplemental File 1, Table 3 for gene family-specific parameters for the above operations)^57^. The resultant ORFs were combined with the initial query file as well as a file containing the protein sequences of the ∼10 closest paralogs to the gene family of interest. This combined fasta was then aligned using MAFFT and a protein phylogeny inferred using FastTree^58,59^. Strongly supported and well-separated clades of orthologous protein sequences were then manually annotated on the tree using FigTree and the corresponding sequences obtained using TREE2FASTA^60,61^. Only those sequences determined to be nested within the gene family of interest were retained.

The above steps, beginning with the tblastn search of the genomes, were repeated in a second and final round of BLAST and phylogenetic clustering^55^. In this round, the sequences obtained from the first round served as queries. Additional rounds would have been performed if new hits had been detected. Orthology and paralogy relationships were again inferred through manual annotation of well-supported clades on the resulting phylogenetic tree. To confirm the robustness of the phylogenetic clustering underlying our annotations, five additional combinations of aligners and tree-building programs were applied to the dataset, generally yielding highly concordant results (Supplemental File 4). For visualizing copy number, we separated gene families into orthogroups likely to have been single copy in the ancestor of the genus *Drosophila*. Finally, we confirmed orthogroup assignments by reciprocal BLAST against the *D. melanogaster* transcriptome, ensuring each sequence matched the expected gene(s). Associated code for the described pipeline for ortholog and paralog identification can be found in Supplementary File 2, a more detailed guide for this methodology and instructions on usage can be found in Supplementary File 3.

#### Inference of gene duplication and loss events via gene tree-species tree reconciliation

To determine whether network genes have undergone correlated changes in copy number across lineages, we sought to infer where on the species tree gene gain and loss events likely occurred. To this end, we applied a method called gene tree-species tree reconciliation using the package GeneRax^62^. The basis of this method is that the relationships among members of a gene family are necessarily linked to the relationships among the species that carry these genes. This is because changes to these genes in copy number and presence/absence represent events that occurred along the branches of the species tree. This means that the pattern of duplication and loss events underlying the complement of genes present in each taxon can be inferred along lineages by “reconciling” the differences between the gene tree and species tree (i.e. making it so that every split in the gene tree corresponds to either a speciation or gene duplication event on the species tree). We applied this method to each orthogroup, here defined as all descendants of genes inferred to be single copy in the common ancestor of *Drosophila*.

We first constructed maximum-likelihood protein phylogenies for each orthogroup (for SP and SPR only, the relevant protein sequences were obtained from Hopkins *et al.* (2024) rather than from our annotations). Protein alignments were constructed using PROMALS3D, an alignment program that integrates structural information into its predicted alignments and has been used to investigate deep protein family relationships^50,63^. Sites with less than 50% occupancy for aligned proteins were then masked with Geneious. The alignment was subsequently manually checked for likely pseudogenes and technical duplicates due to assembly errors, as described below, and offending sequences were removed from the alignment. The starting gene tree was then inferred from the resulting alignment using RAxML-ng, and the starting species tree, taken from Kim et al. (2024), was trimmed to remove species not present in both our analysis and Hopkins *et al.* (2024) (leaving 128 spp.)^24,45,53^. These files, alongside the inferred protein alignment, were used as the inputs for GeneRax and will be made available in a data repository upon publication. In all cases, only duplications and losses (i.e., no horizontal transfers) were allowed (see Supplemental File 1, Table 10 for all relevant parameters).

#### Ruling out spurious duplication/loss events due to assembly errors

To rule out assembly errors as a major driver of the patterns observed, we manually validated a subset of the inferred gain and loss events. For gains, to rule out technical duplication due to (for example) incorrect haplotype phasing, we manually checked all inferred gene duplications on terminal branches (i.e., those observed only in a single genome). We flagged and removed any cases where the duplicates are both 1) identical in DNA and amino acid sequence, and 2) not convincingly shown to represent unique sites in the genome (e.g. the two copies are in tandem on single scaffold/contig). Across all gene families, we removed 3 probable technical duplicates from the dataset and adjusted relevant analyses accordingly. For losses, validation was performed only for those events associated with the four major concerted losses of sex peptide network members that were identified prior to the calculation of correlations. To do this, for long-read assemblies, we confirmed losses syntenically by checking the presence/order of conserved genes flanking the gene that was lost (Supplemental File 1, Table 12). To identify locations of flanking genes, we queried *D. melanogaster* protein sequences against other species’ genomes using tblastn. Alternatively, for species with only short-read assemblies, as is the case for many *montium* group species, we searched for evidence of the lost genes in the raw sequence reads to rule out assembly artifacts. We used the amino acid sequences of each missing gene as a query in the tblastn searches against the raw reads associated with each assembly. We did this for the species with losses as well as for several close outgroups that did not experience the loss events of interest and compared the distribution of e-values obtained for each query between these groups (Supplemental File 1, Table 9). The expectation is that, in the case of a genuine loss (either pseudogenization and degeneration or total loss), the e-values of the hits obtained should decay much faster than for closely related species where the gene is intact. The only exception to this would be very recent pseudogenes that have degenerated minimally. In these cases, we used the assembly to confirm the presence of a frameshift mutation or premature stop codon in the coding sequence. We also blasted the best hit obtained back to the *D. melanogaster* transcriptome to screen for the presence of pseudogenes. No spurious losses were identified.

#### Correlation of gene duplication and loss among orthogroups

Statistics were computed in R v. 4.5.1^64^. To test for correlated changes in copy number among gene families, we performed Spearman’s rank correlations on the raw counts of events (i.e., gains or losses) inferred to have occurred on each branch of the species tree. This approach is conservative in that it identifies only cases where two gene families experience events of the same kind on the same branch, meaning that some types of genuine associations may be overlooked. For example, we do not test for associations between events of different types (e.g., when losses of orthogroup A are coupled to gains of orthogroup B) or across time (e.g., when ancestral gains of orthogroup A drive subsequent gains of orthogroup B on descendent branches). Also note that, for identifying correlated loss events, only complete losses of all members of orthogroups (as defined above) were considered. There are two reasons for this: 1) duplication followed quickly by sporadic loss of a single daughter copy in descendent lineages likely carries different effects than complete loss of an orthogroup that predates the genus, and 2) because we don’t allow for gene transfer in our models, introgression or horizontal gene transfer could be misinterpreted as an ancient duplications followed by multiple subsequent loss events, thereby adding considerable noise to the analyses.

#### Testing whether a candidate gene functions in the Sex Peptide Network

Fly stocks used in this study are listed in Table 1 (in Supplementary File 5). Control and experimental flies were maintained on a standard medium containing sugar, oatmeal, cornmeal, agar, and yeast, under a 12h light:12h dark cycle at 25°C. Flies were collected after eclosion and aged for 4-7 days in single-sex groups. Males were individually aged in a flat bottom 1.5 mL 96 well block filled with 0.5 mL food per well and covered with a PCR foil with air holes. Unmated females were aged in groups of 10-15 in food vials. All females used were from the wild type Canton-S strain. The detailed genotypes of experimental male flies are listed in Table 2 (in Supplementary File 5), but are briefly listed here as the two independent knockdown lines (*tub-GAL4 > RNAi BDSC_65910* and *tub-GAL4 > RNAi VDRC_14158*, with associated controls *RNAi BDSC_65910* control and *RNAi VDRC_14158* control, respectively; *tub-GAL4* flies were used as an additional control).

For remating assays, individual couples of 1 experimental or control male and 1 wild type Canton-S virgin female were observed in mating chambers (1 cm diameter, 4 mm height). Females that copulated within 1hr were recollected after the end of copulation and kept (without males) for 4 days. The mated females were then individually tested for remating with wild type Canton-S males within 1hr in a mating chamber.

## Results and Discussion

Between species, changes in gene copy number of Sfps caused by duplication or loss events could arise through neutral or adaptive processes. Copy number of Sfp genes may increase due to strictly neutral processes, since Sfps, as secreted proteins, are likely to be dosage-tolerant, and paralogs may show functional redundancy which could buffer against gene loss^4,65–67^. Alternatively, copy number increases may be driven by natural selection to rapidly adjust dosage in response to selective pressures, and duplication can provide the evolutionary flexibility to allow a paralog to explore new functions without compromising the original gene copy^36,68–73^. Similarly, gene loss of Sfps may also be caused by a neutral process such as loss of selective constraint through redundancy or decreases in functional importance^36,69–71^. Gene loss may also occur as an adaptive process, such as when maintaining the gene imposes fitness costs (e.g. for reproductive proteins, through selection, or loss of benefit under changed mating systems or ecology)^74,75^. However, correlations in gene duplication or loss events between functionally related genes could point to whether directional selection informs CNV^76–78^. Here, we investigate the origin and CNV of a network of Sfps (and some functionally related female proteins) that work with the Sfp Sex Peptide (SP) to modulate the long-term post-mating response in *Drosophila* females. To exhaustively identify all homologs and accurately distinguish between paralogy and orthology, we developed a pipeline pairing iterative queries of *Drosophila* genomes with phylogenetic clustering of hits to resolve accurate orthologous vs paralogous relationships. Statistical tests revealed covariation in gains and losses among certain functionally related Sfp proteins, suggesting that functional interactions between these proteins may underlie covariation, and that the observed CNV of Sfps is unlikely to be explained solely by neutral evolutionary processes.

### The origin of Sex Peptide Network proteins largely predates Sex Peptide’s origin

Of the eight known members of the SPN Sfps, four are predicted active or inactive serine proteases, two are lectins, and two are CAP domain proteins^35^. For protein families that contribute to the SPN, we constructed protein phylogenies of all *D. melanogaster* members. We determined that all SPN proteins belonging to the same protein family were closely paralogous to each other, and frequently have additional closely paralogous Sfps that have not yet been tested for network function^35^ (Figure 1B). The close paralogy between many network members suggests that duplication followed by specialization of function may have contributed to the origin of the network^70^. We tested this by an iterative BLAST approach across 135 genomes, including species within the genus *Drosophila* as well as outgroup species beyond the genus^55^. This approach not only allowed for a detailed profiling of protein evolution over ∼40 million years, but also increased the likelihood of identifying orthologous proteins that may exhibit elevated sequence divergence^24^. By iterating through progressively more distantly related species, the method enhanced sensitivity and helped avoid missing genes that could be obscured by piecemeal sampling within the genus. Notably, *seminase*’s frequent and complex pattern of duplication and loss between species led to many distinct lineages of *seminase* homologs across the genus. To trace changes in CNV across the genus, we combined all *seminase* paralogs into a single orthogroup, composed of the set of genes across species that are descended from a single ancestral gene (here, ancestral to the genus *Drosophila*).

We traced the origins of SPN genes (Figure 2, Supplementary File 1, Table 14). Our findings show that all members of the SPN are present in the common ancestor of the genus *Drosophila*. Additionally, all the network members are identifiable in the outgroup Drosophilinae fly species *Chymomyza costata* and *Scaptodrosophila lebanonensis*, suggesting that orthologs of all known network members existed in the ancestor of the Drosophilinae within the Drosophilidae family of flies. SP itself also originated in the ancestor of Drosophilinae^12,45^. However, many network members are detectable in the Drosophilidae subfamily Steganinae species *Leucophenga varia* (including *Sems*, *CG9997*, *Aqrs*, *CG17575*, *Antr*, and *CG1652*). These members represent all the paralogous groups of network proteins described in Figure 1A and include all network members required for transfer of other network proteins. This suggests that, except for duplication of *CG9997* and *CG1656* (to give rise to *intr* and *CG1652* respectively), the network originated in the common ancestor of *Drosophilidae*, predating the origin of SP. Notably, no orthologs of these proteins—or their close paralogs, which are also seminal fluid proteins in *D. melanogaster*—can be detected outside of *Drosophilidae*. Taken together, these findings may suggest the network existed before the emergence of SP with its own independent function, and that SP only later co-opted the network for its own function.

**Figure 2:**
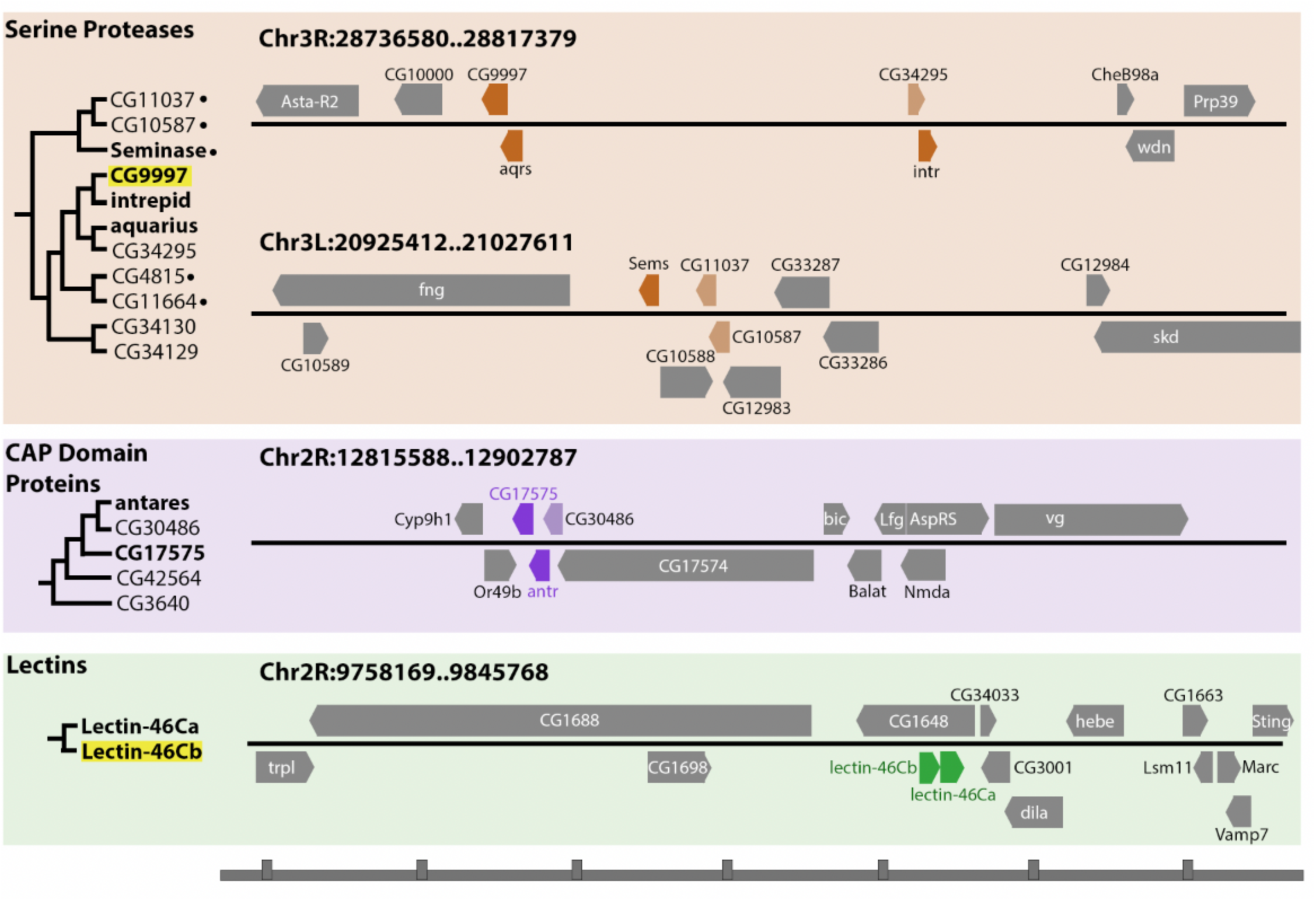
Sex Peptide Network seminal fluid proteins arose through gene-duplication. SPN proteins are closely paralogous to other Sfps including some that are known members of the SPN. Protein trees are derived from the phylogenetic clustering described in the Methods to establish orthology and paralogous relationships and supported by >50 bootstraps. All SPN proteins are bolded and they, along with their paralogs, are all known Sfps^15^. Circles to the right of serine proteases protein names indicate that they are predicted to be active proteases, others are annotated as inactive proteases. Highlighted genes have been shown to evolve under positive selection^46^. Genomic regions containing SPN proteins in *D. melanogaster* (r6.65) are depicted^99^. Tandem duplication appears to contribute to the origin of these Sfps. Darker colored boxes are SPN members, lighter colored boxes are paralogous Sfps whose SPN function is not known. ORFs and distances are approximate and are not shown perfectly to scale. Ticks on x-axis represent ∼12.5 kb.

### Extensive copy number variation of Seminal Fluid Proteins in the Sex Peptide Network

The presence of all SPN orthogroups in the *Drosophila* genus ancestor enables a 40-million-year comparative analysis of gene duplication and loss. By comprehensively annotating orthologs of SPN proteins and their close Sfp paralogs, we uncovered substantial CNV among many Sfps, arising from both gene gain and loss events (Figure 3).

**Figure 3.**
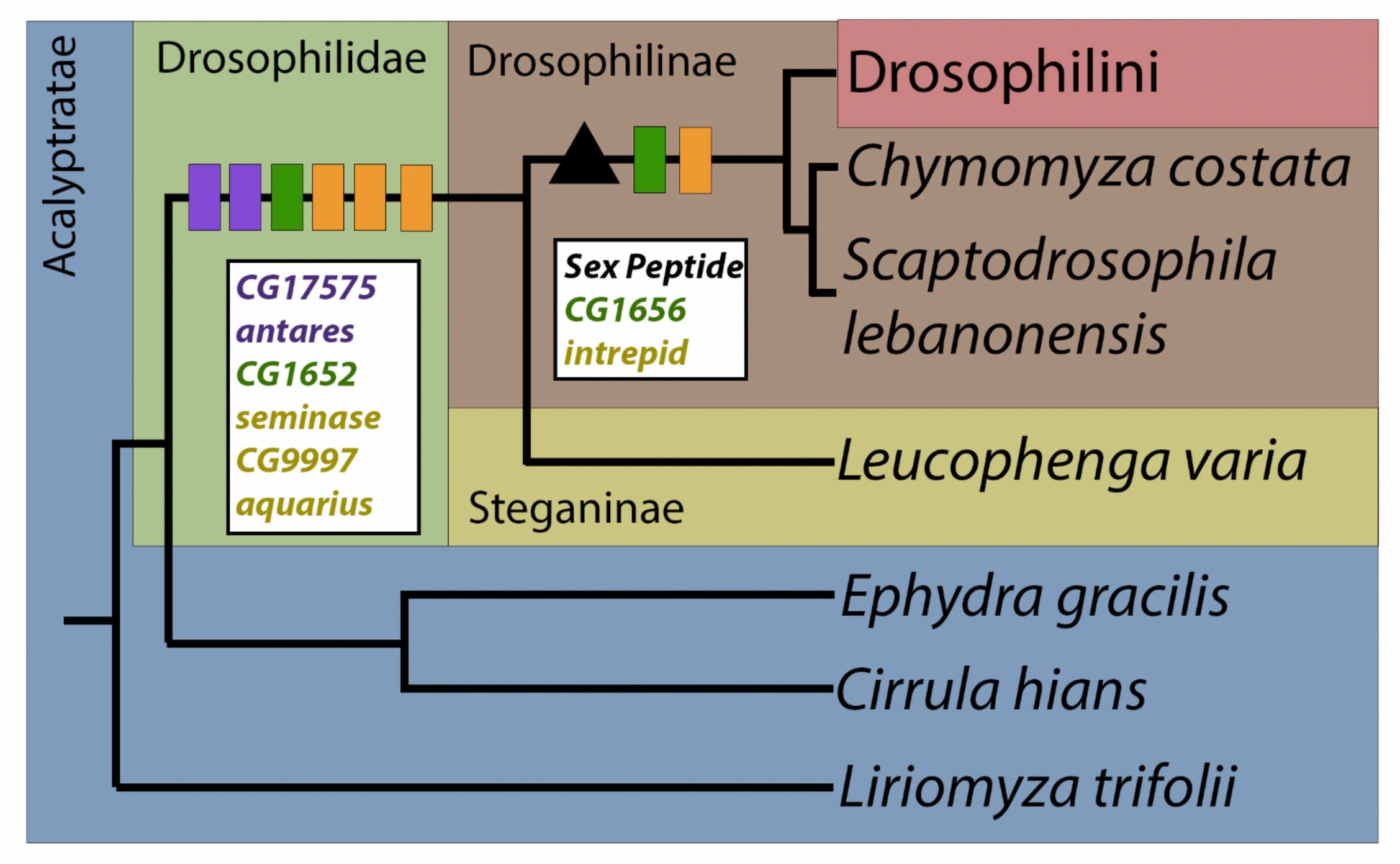
Many SPN Sfp genes arose before the Sex Peptide gene. Many of the SPN Sfp genes (*CG17575*, *antares*, *CG1652*, *seminase*, *aquarius* and *CG9997*) arose in *Drosophilidae*, and predated the origin of SP in *Drosophilinae*^12,45^. Along the lineage where SP arose, a duplication of *CG1652* and a duplication event of *CG9997* led to the origin of the two other known members of the SPN (*CG1656* and *intrepid* respectively).

**Figure 4.**
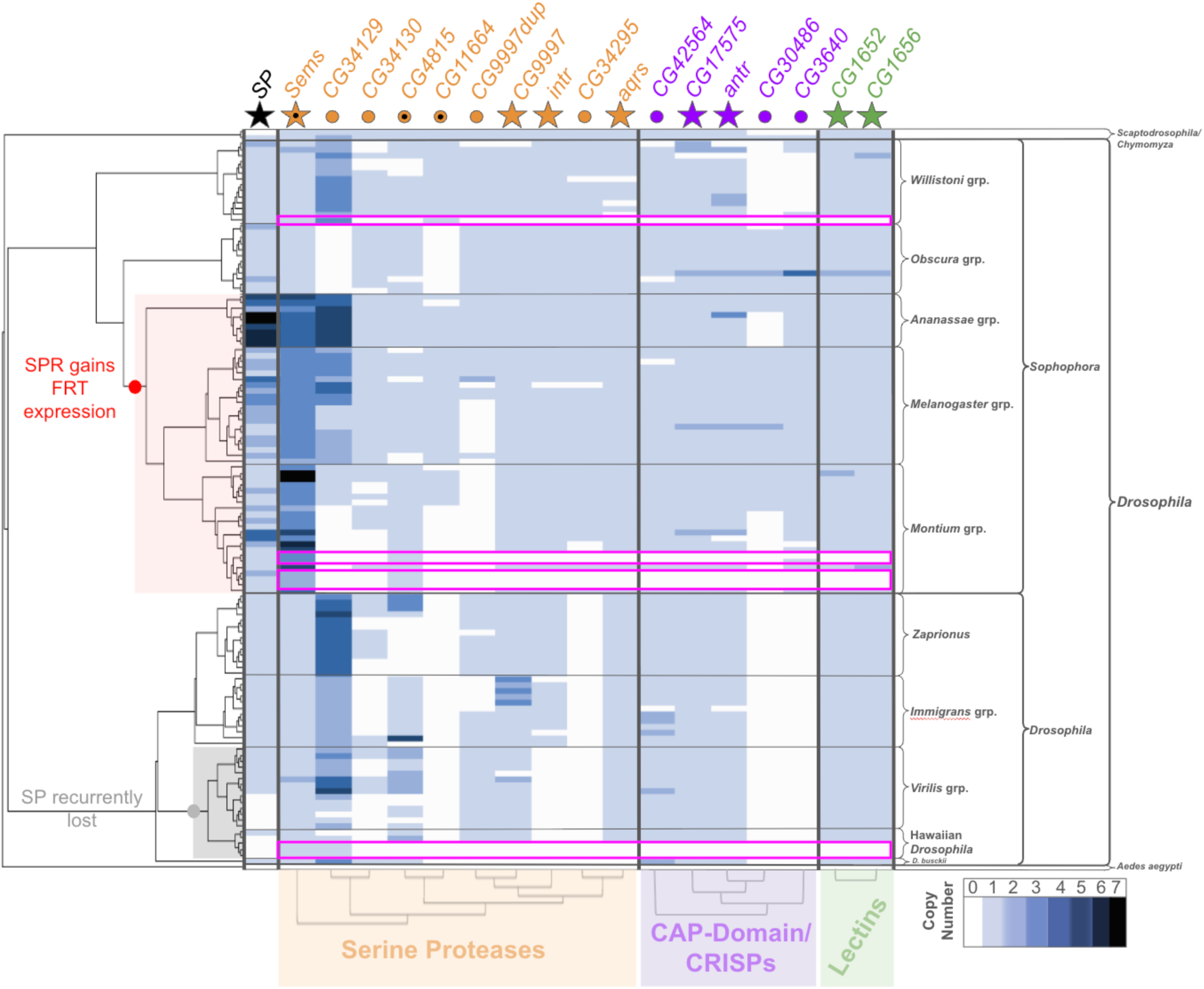
Copy number variation of SPN seminal fluid proteins in the genus *Drosophila*. Copy numbers of SPN genes and their close paralogs for 135 Drosophilids inferred via iterative blast and phylogenetic clustering (except *SP*, for which copy number data were obtained from Hopkins *et al.* (2024)). Genes are colored by protein type; Black = Sex Peptide; Orange = Serine Proteases; Purple = CAP-domain genes/CRISPs; Green = Lectins. Stars denote genes with known SPN function in *D. melanogaster*. For serine proteases, those stars/circles marked with a black dot denote active proteases, whereas unmarked shapes represent inactive protease homologs. The four main concerted losses of most network genes we identified are bordered by pink boxes. The point on the species tree where SPR was inferred to have gained expression in the female reproductive tract is marked in red^47^. Similarly, lineages where *SP* is recurrently lost are marked in gray. CNV data underlying the figure can be found in Supplementary File 1, Table 4.

**Figure 5.**
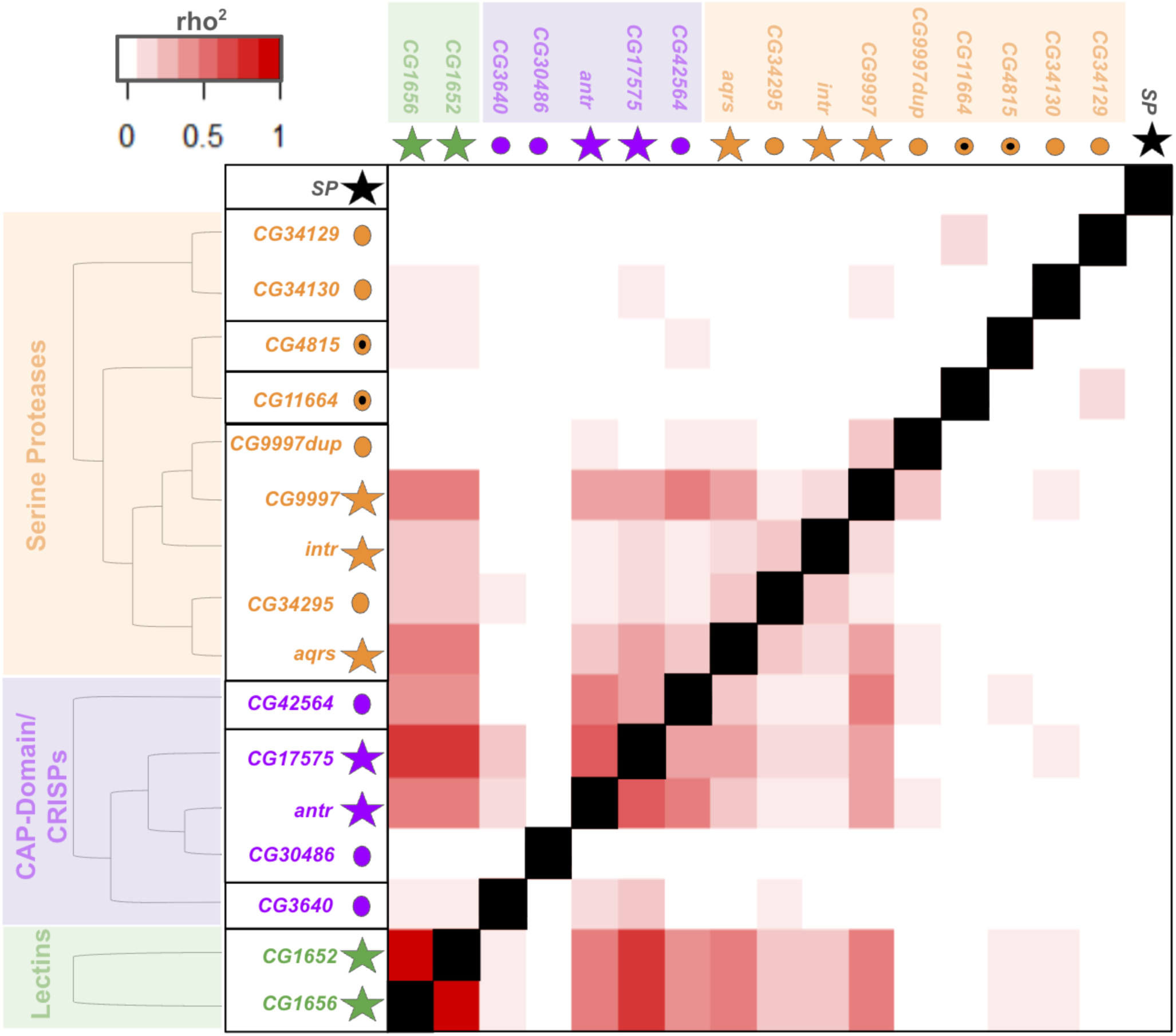
Correlated loss events between members of the SPN. Pairwise Spearman’s rank correlation coefficients (rho; squared for visualization purposes) between SPN genes and their close paralogs for concerted total orthogroup losses by branch (associated values can be found in Supplementary File 1, Table 7). The location of loss events on the species tree was inferred by gene tree-species tree reconciliation. For the correlations, each branch (n = 258) was assigned a value (0 or 1) denoting whether or not a loss occurred on that branch, and the resulting vectors for each orthogroup were correlated. Genes are colored by protein type; Black = Sex Peptide; Orange = Serine Proteases; Purple = CAP-domain genes/CRISPs; Green = Lectins. Stars denote genes with known SPN function in *D. melanogaster* prior to this study. For serine proteases, those stars/circles marked with a black dot denote active proteases, whereas unmarked shapes represent inactive protease homologs.

Against our expectations, aside from *seminase* and CG43129, no SPN genes and paralogous Sfp genes investigated exhibit the exaggerated CNV observed for SP, indicating that functional constraints limited the expansion of these SPN members. Still, small numbers of duplications (∼1–8) occur sporadically for nearly every gene. Loss events are even more striking. In 9 species spanning the genus the entire SPN is not present, with the exception of *seminase* which is never lost. This recurrent, wholesale loss is unexpected given the ancestral origins of SPN genes and their crucial role in long-term post-mating responses in *D. melanogaster*, although their function in other species is not known. The pattern suggests functional interdependence among these network members such that the loss of one renders others dispensable. Notably, *seminase* and usually *SP* persist in these lineages, implying that their retention may reflect functions independent of the rest of the network (e.g. *seminase*’s interaction with *Semp1*^79,80^).

The SPN gene *seminase* as well as *CG34129*, a Sfp paralog of the serine protease SPN genes show extensive CNV across the genus driven by duplication events. SP shows 34 independent duplication events, and *seminase* 28, generating complex patterns of lineage-specific homology. In *D. melanogaster*, *seminase* has two close tandem paralogs (*CG11037* and *CG10587*), neither of which has yet to be functionally tested for SPN functions. Duplications of *SP* and *seminase*, but not *CG34129*, are largely restricted to clades where *SPR* is expressed in the female reproductive tract (FRT)^45–47^.

CNV patterns provide insights into functional relationships and selective pressures. Duplications may be retained neutrally, by selection for increased dosage, or through neofunctionalization. Comparative –omics analyses can help distinguish these scenarios: divergence in sequence or expression suggests gains in copy number are driven by selection for novel function or correlated dosage-dependency. However, most duplication events of investigated Sfp genes are recent and specific to a few species, potentially implying that duplication events are not rare but short-lived, indicative of neutral evolution or frequently fluctuating selective pressures between species. Identifying correlations of copy number changes with other functionally related genes would reject neutrality in favor of selection. However, only *SP*, *seminase*, and *CG34129* display sufficient CNV for such tests, as most SPN Sfps appear resistant to duplication. Meanwhile, loss events can be seen as an inevitable outcome when a gene’s function is no longer needed or perhaps when the gene has become deleterious. The repeated, coordinated losses of SPN members highlight a paradox: genes deeply conserved and essential in some species can become dispensable in others. This turnover underscores the fluidity of reproductive networks and suggests that evolutionary pressures on mating systems can rapidly restructure even highly integrated molecular networks. The repeated occurrence of lineages where all SPN members except *seminase* are lost could suggest interdependency in function between most of the SPN genes such that loss of one leads to the loss of functionality of the others. Interestingly, in most of these lineages where most SPN proteins are lost, *SP* is still present, indicating that these shared losses may be independent of *SP* function.

### Correlation of gene duplication events of *Sex Peptide* and *Seminase*

Frequent duplication of *SP*, the SPN gene *seminase*, and the Sfp *CG34129* across the genus *Drosophila* provided an opportunity to test whether duplication events among these genes are correlated. Rare duplications cause meaningful correlation analysis to be infeasible since technical noise could overshadow genuine signals. When a gene is duplicated and fixed in a species, that duplication will generally be inherited by descendant species, which may create correlations in copy number across lineages simply due to shared evolutionary history. To avoid the effects of phylogenetic nonindependence confounding our results, we tested for co-occurrence of discrete duplication events along branches of the species tree rather than raw copy number covariance (Supplementary File 1, Table 5 shows number of duplications along branches). Such correlations may indicate shared selective pressures driving increased copy number (e.g. pressures favoring increased dosage or evolutionary flexibility), or possibly some functional relationship between protein products where duplication of one gene promotes duplication of another^78,81–85^.

Our analyses revealed a striking pattern: duplication events of *SP* and *seminase* are highly correlated across *Drosophila* lineages (Spearman’s correlation, p = 0.003, ρ = 0.19), whereas no such correlation was observed between *SP* and *CG34129* (p = 0.3, ρ = .064), or between *seminase* and *CG34129* (p = 0.2, ρ = .089). Specificity in correlation between *SP* and the SPN member *seminase* but not the paralog *CG34129* suggests that frequent duplication of *SP* and *seminase* is evolutionary related and not likely driven by neutral evolution. Further, both *SP* and *seminase* show an elevated rate of duplication in lineages where *SPR* gained expression in the FRT (*SP*: 4.0 vs. 0.22, 28 duplications over 7 units of branch length in the *SPR* in FRT clade vs 3 duplications in 13.5 units of branch length; *seminase*: 2.7 vs. 0.30, 19 duplications over 7 units of branch length in the *SPR* in FRT clade vs 4 duplications in 13.5 units of branch length duplications per unit branch length). We tested whether duplication events of *SP* and *seminase* were more common along branches in the clade where *SPR* has acquired expression in the FRT. We used a chi-squared test to compare the observed occurrences to expected values based on the total number of events observed and the total branch length for branches in the clade with SPR expression in the FRT or branches outside this clade (Supplemental File 1, Table 11). We observed that *SP* and *seminase* were more often duplicating in the clade where SPR gained FRT expression (*SP: p* <0.0001, *seminase: p* <0.001). No other genes with duplications shared this pattern. This indicates that after the origin of the *SPR* FRT expression, strong selective pressure acted on both *SP* and *seminase* to expand in copy number. In contrast, while Seminase acts in a cascade to cleave Semp1 (which then processes Acp36DE and ovulin), CNV of *Semp1* is not correlated with *seminase* or *SP* Supplemental Figure 2. This implies that *seminase*’s extensive duplication may be tied to its interaction with *SP* or to something else that also drives *SP*’s duplication rather than to its broader protease cascade function.

For Sfps with identifiable homologs across species, copy number can vary dramatically. The repeated co-duplication of *SP* and *seminase* cannot be explained by neutral processes alone, instead suggesting selective pressures or interdependence drive their CNV across species. Whether this correlation reflects unidirectional dependence (e.g., *SP* expansions occur following *seminase* duplication) or shared selective pressures acting on both genes remain unresolved, but either possibility points to a genuine functional connection. The increased rate of duplication of these orthogroups following SPR’s acquisition of FRT expression further supports the idea that this transition amplified selective pressures on *SP* and *seminase*. Yet this raises a key question: why do *SP* and *seminase* display co-duplication, but not with the other identified SPN genes? One possibility is that *seminase* is unusual within the network: although required for *SP* and other Sfps to bind sperm, *seminase* itself is not detectably associated with sperm as has been seen for most of the other Sfps. This difference may make *seminase* more tolerant to duplication, and therefore more likely to co-expand with *SP*, whereas sperm-associated proteins may have to remain more constrained. In summary, the correlated duplication of *SP* and *seminase* suggests a potential selective and functional dependency, highlighting the value of phylogeny-based duplication analyses in uncovering unexpected gene interactions, and in generating specific, testable hypotheses about the molecular mechanisms driving the correlations.

#### Correlation of gene loss events between members of the Sex Peptide Network, but not with Sex Peptide

We also investigated correlations in gene loss events between SPN proteins across the genus *Drosophila* as an alternative signal of variation in natural selection acting across species. For our analysis we only considered complete losses of all copies within an orthogroup ancestral to the genus *Drosophila*, rather than individual loss events of a subset of paralogs (Supplemental File 1, Table 6 (orthogroup absence) & Table 7 (Spearman’s rho).

We observed strongly correlated losses between all members of the network except *SP* and *seminase* (*SP* is lost 3 times, but the losses are uncorrelated). Indeed, signatures of correlated loss between SPN members appear to be stronger than the reported ERC between the same network member pairings. These effects are largely driven by branches upon which these SPN Sfps are lost at once. In three cases, all SPN genes (except *seminase*) disappear simultaneously along single branches (out of 258 total branches on the tree and less than 10 total losses per gene); in a fourth instance, *intr* is lost first, followed by the remainder of the network on a descending branch (lagging events would not increase statistical significance with our approach). We investigated the causal event of the losses of SPN proteins in the four lineages that lost all network members and, in many cases, decayed ORFs of pseudogenized genes were identifiable, indicating independent gene loss events rather than loss of multiple tandem genes via deletion of genomic regions, enabling resolution of coordinated loss events even between genomically adjacent paralogs.

Although pairwise comparisons between most network members showed statistically significant correlations among loss events, the biological meaning or strength of these correlations remains unclear. However, it is intriguing that the strongest correlation in loss events for *CG1652* and *CG1656* is to the three network Sfps that are required for their successful transfer (*Aqrs*, *Antr*, *CG9997*), suggesting functional interdependence driving their strongly correlated losses. Potentially, this could indicate physical interactions in a complex where loss of one protein leads to the destabilization of the complex and the loss of the other members, as was seen for correlated losses of PRDM9 complex members in mammals and for complexes in other organisms^78,81,82^. The disconnect in correlated loss events of *seminase* and *SP* to the rest of the SPN may be related to these genes’ function. Since the SPN genes largely arose before *SP*, potentially these genes have an ancestral shared function independent of *SP*.

#### Sex Peptide network gene loss correlates primarily with loss of other known Sex Peptide Network genes

The coordinated losses of SPN Sfps observed is suggestive of selective forces that act upon the network as a whole. However, this pattern is also consistent with pressures acting on Sfps more broadly to cause loss events along specific lineages, leading to strong signals of coordinated CNV independent of pressures on the network *per se*. If coordinated losses reflect lineage-specific selective pressure on Sfps and are not specific to SPN members, then we would expect Sfps without association with the SPN to not show correlation of loss events with SPN members. Indeed, not all Sfps paralogous to SPN members showed correlated patterns of loss, indicating that this signature is more likely to be specific to network members. *CG9997-dup* (absent in *D. melanogaster* but ancestral to genus *Drosophila*) is a close tandem paralog of SPN members but does not show correlated losses with any known SPN members. Also, *CG30486*, a close tandem paralog of *antares*, shows no correlation of loss events nor ERC. Previous genetic investigation of this gene showed no role in the SPN.

To more strongly query our hypothesis that correlation in loss was specific to proteins that are part of the SPN, we additionally investigated 30 other Sfps for shared patterns of loss with SPN proteins (Supplementary File 1, Table 13). We chose Sfps belonging to the same protein families as known SPN members but which are not closely related to any of them, and Sfps representing other protein families. We investigated whether any of these Sfps show losses along the same four lineages where all known SPN members are lost. We used reciprocal BLAST to screen for the presence/absence of these Sfps within genomes of the species where all network members were lost as well as a few closely related taxa where they remained intact. None of the examined Sfps showed losses along lineages in which all network genes were lost. This result suggests that the detected correlation in loss events between members of the network reflects their shared roles, rather than overall pressure on a similar category of proteins. The specificity of correlation of losses to members of the SPN or their paralogs (with the exception of *SP* and *seminase*), and not Sfps more broadly points to the usefulness of this metric in identifying functionally related proteins in a guilt by association-type approach. However, since the method requires gene loss, it can only be applied to orthogroups that are dispensable in some lineages and ancestral enough for repeated loss events to occur.

#### New SPN members can be identified as genes whose losses correlated with Sex Peptide Network Genes’

Interestingly, correlations in loss events were also observed between SPN members and some paralogous proteins whose SPN functions were unknown. Further, these Sfps showed correlated losses with each other. A striking example of this is the Sfp *CG42564*, which shows correlated losses with the SPN members to which it is closely paralogous, as well as to other unrelated SPN members. This result either indicates correlation in loss events unrelated to SPN function, perhaps related to its close evolutionary relationship to the SPN proteins or shared selective pressure acting broadly on Sfps, or could suggest that the *CG42564* gene is an uncharacterized member of the SPN. Supporting the latter hypothesis, in all four lineages where all known SPN members are lost, *CG42564* is also absent. To test whether *CG42564* is, in fact, a new member of the SPN, we assayed a long-term post-mating response in mates of males knocked down for *CG42564*, using two independent RNAi lines, and comparison to controls. In both independent knockdowns tested, mates of knockdown males regained receptivity to mating, in contrast to mates of control males (remating at 4 days; for line BDSC 65910 – 36/49 females (73.5%) mated to knockdown males vs 0/42 females mated to control males, for line VDRC 14158 – 46/54 females (85.2%) mated to KD males vs 0/23 females mated to control males; similarly, 0/48 mates of *tub-GAL4* males re-mated; Fisher’s exact test for all knockdown/control remating comparisons: p<0.0001), indicating that *CG42564* is indeed a member of the SPN. Functional investigation of other genes that show correlated loss with SPN genes may reveal additional new members of the SPN.

#### Female genes that modulate Sex Peptide function downstream of SPN action do not show extensive CNV or correlation of gene duplications or losses with Sex Peptide or its network

The SPN Sfps transferred by the male during copulation are essential to SP’s binding to sperm and long-term effects on the female, however, female factors are involved in mediating SP’s downstream effects on female post-mating physiology and behavior independent of the SPN^13,35,44^. Most famously in *Drosophila melanogaster*, SPR (Sex Peptide Receptor) mediates many of SP’s effects on the female^86^. However, despite their crucial interaction, there is no correlation in gene CNV between *SP* and *SPR*, with *SPR* consistently found as a single-copy gene in all species where it has been identified, and no correlation in sequence evolution of these two genes is seen by ERC^45^.

Here we investigated CNV of three additional female-expressed genes that mediate downstream effects of SP in females (*Frma*, *Hdly*, and *Esp)* which, unlike *SPR*, were identified via correlated sequence evolution with *SP* via ERC^13^ (Supplementary Figure 3). *Frma* is a neprilysin-like metalloendopeptidase, a class of proteins common among seminal fluid proteins; *Esp* is a member of the SLC26 family of anion exchangers; *Hdly* is an orphan gene with no known structural domains^49^. *SPR*, *Hdly*, and *Esp* originated prior to the emergence of SP and its associated network, with clear one-to-one orthologs identified in *Aedes aegypti*, the most distantly related outgroup examined, *Frma*’s origin appears to predate the common ancestor of the examined taxa (Supplementary File 1, Table 14). This suggests that while their specific functions and expression in the FRT across species are unknown, most of these genes are ancestral to Diptera, predating SP and the SPN. No significant CNV was observed for any of these genes across the genus, although *Hdly* and *Esp* were occasionally lost in some lineages. Therefore, as was the case for SPR, there is no CNV correlation between these genes with each other, *SP*, or the SPN.

While it is noteworthy that female proteins that mediate SP’s downstream effects in *D. melanogaster* show uncorrelated CNV with *SP* and its network, this pattern may not be unexpected due to broader functional constraints impacting these genes’ evolution. SPR, for example, has an alternative ligand and was likely co-opted as the SP receptor within the genus, limiting the extent of shared evolutionary pressures with SP^87^. Other female modulators may be similarly constrained: protein properties such as expression specificity, molecular role, or network connectivity can strongly influence tolerance to CNV^77^. Meanwhile, secreted proteins such as Sfps may be less constrained in CNV due to greater functional flexibility and lower structural constraints compared to transmembrane or signaling proteins like SPR and Esp^29–31^. Although both male and female reproductive genes have been shown to diversify rapidly between species, sometimes driven by positive selection, this pattern is generally observed to be more pronounced in male-expressed genes^1,4,88^. However, differences in gene category and function are frequently overlooked as contributors in these comparisons.

## Conclusion

Lineage-specific evolutionary pressures –whether positive, negative, or relaxed selection– affect genes in ways that shape the evolution of their sequence, expression levels, and copy numbers. These effects leave detectable signatures that can be analyzed to generate testable hypotheses about underlying molecular mechanisms. Phylogenetic profiling methods harness such patterns to infer functional linkages between traits or molecular partners subject to shared evolutionary pressures^78,89–91^. Most famously among genomic approaches, Evolutionary Rate Covariation (ERC) identifies functionally related genes through correlated sequence changes along phylogenetic branches and has been successfully applied to diverse biological processes, including in *Drosophila*^13,14,92–95^. Indeed, many SPN proteins in our study were first identified through ERC^13^. Recent increases in available genome sequences spanning deep evolutionary timescales have expanded opportunities to use patterns of gene loss and gain to identify conserved members of protein complexes^23,25^. Here, by combining newly available, high-quality *Drosophila* genome sequences with a high-throughput approach for orthology and paralogy annotation via phylogenetic clustering, we characterized copy number changes within a recently evolved protein network that shows recurrent duplication and positive selection. Our detection of correlated loss events among SPN members validates this approach for uncovering functionally related proteins, and reveals one candidate, *CG42564,* as a new SPN member. Correlated evolutionary patterns in sequence, expression, copy number, and other phylogenetic profiling metrics are likely complementary, producing both overlapping and distinct sets of candidate genes. The nature of a gene’s functional relationships may influence which correlations are detectable by different methods. For example, ERC can reflect direct or indirect interactions, whereas shared loss correlations (such as those we observed among SPN members) may primarily indicate more direct interaction, perhaps as members of a complex or direct pathway. ERC also detected downstream female modulators of SP function that were missed by shared loss analysis, consistent with pleiotropy and indirect roles. On the other hand, ERC may struggle to identify correlated patterns of evolution when complex paralogy and orthology relationships (common among the proteins we investigated) are present, although new strategies may mitigate this limitation^96^. Thus, combining ERC with duplication/loss correlations improves candidate identification: overlaps may highlight strong partners, while ERC-only hits may reflect indirect interactions. Both approaches, however, depend on conserved relationships across deep time and are limited by tolerance of duplications, multifunctionality, and lineage-specific copy number dynamics.

## Data Availability

All supplemental tables described in the text can be found in Supplemental File 1. This file contains, among other supporting information, lists of publicly available transcriptomes and genomes used in the analysis, parameters for the sequence collection pipeline, and data underlying figures and statistical analysis. Supplemental File 2 and 3 contain the code used for identifying orthologs and a more detailed description of its usage respectively. Supplemental Figures can be found in File 4. We have also uploaded intermediate analysis files to a data repository upon publication.

## Acknowledgments

We thank Dr. Elissa Cosgrove with assistance in the bioinformatic analysis. We thank Drs. Nathan Clark and Matt Pennell for helpful discussions. Finally, we thank members of the Wolfner and Clark labs for helpful discussions and suggestions.

---

**Supplemental Figure 1:**
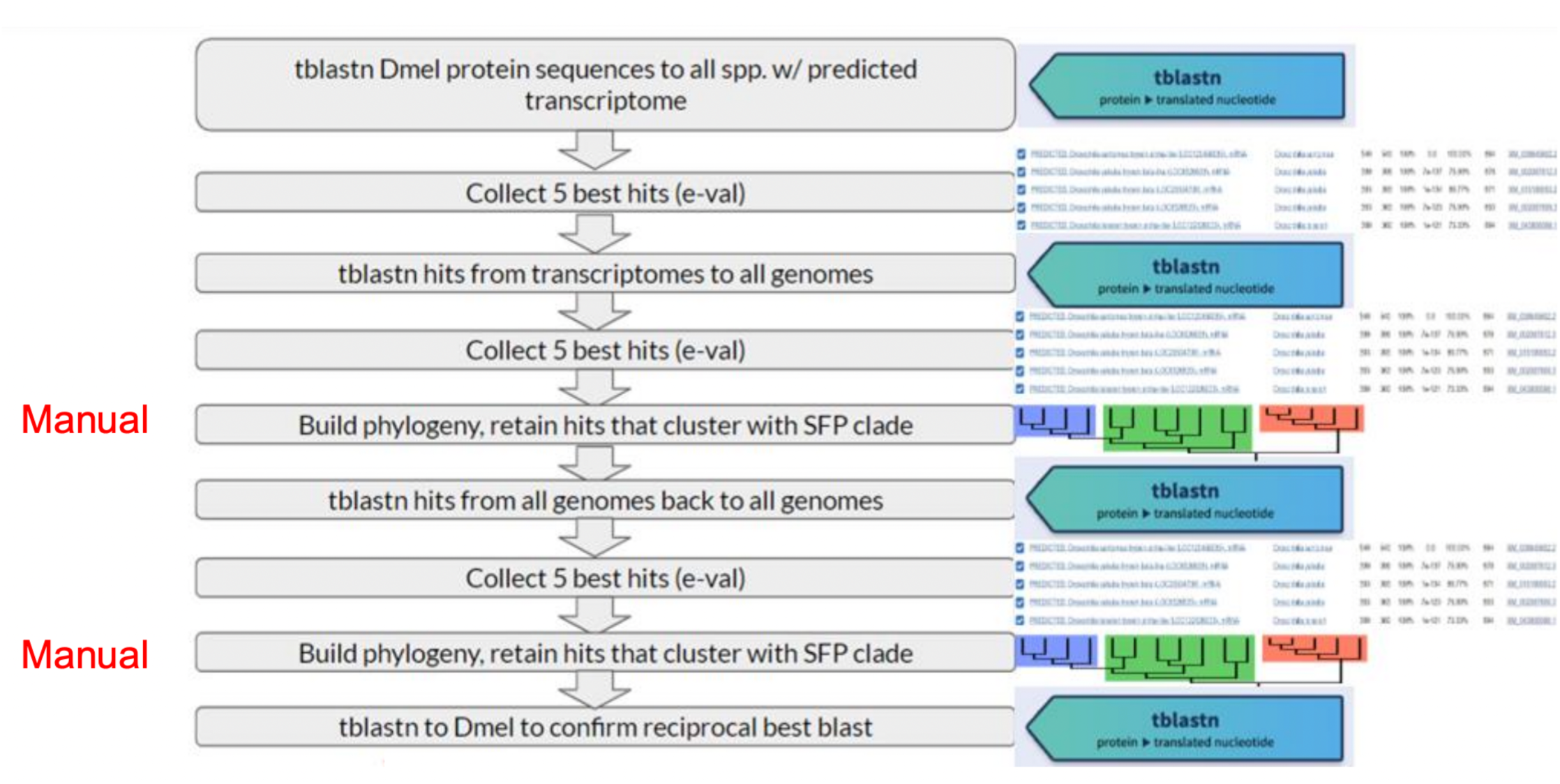
Graphical overview of computational pipeline used for sequence collection and determination of orthology/paralogy relationships via iterative blast and phylogenetic clustering. See Materials and Methods for a more detailed description of the approaches used. Associated code can be found in Supplementary File 2 and additional instruction on implementation in Supplementary File 3. Briefly, *D. melanogaster* protein sequences representing all orthogroups in a protein family of interest were used as the query of a tblastn search to the predicted transcriptomes of 35 *Drosophila* species. The five best hits were then collected for each query protein. The corresponding amino acid sequences were subsequently obtained from NCBI and themselves used as the query for a new tblastn search, this time to the genomes of 135 Drosophilids. The five best hits for each query were translated, aligned, and used to build a protein tree containing the hits recovered from all species. The hits that convincingly clustered with the gene family of interest on this tree used as the query of a final tblastn search of the same 135 genomes. The subsequent filtering and phylogenetic clustering were repeated, and final orthogroup designations manually annotated on the resulting tree. To additional validate our annotations, we confirmed that each protein sequence, when used as the query of a tblastn search of the *D. melanogaster* predicted transcriptome, hits the expected *D. melanogaster* transcript.

**Supplementary Figure 2.**
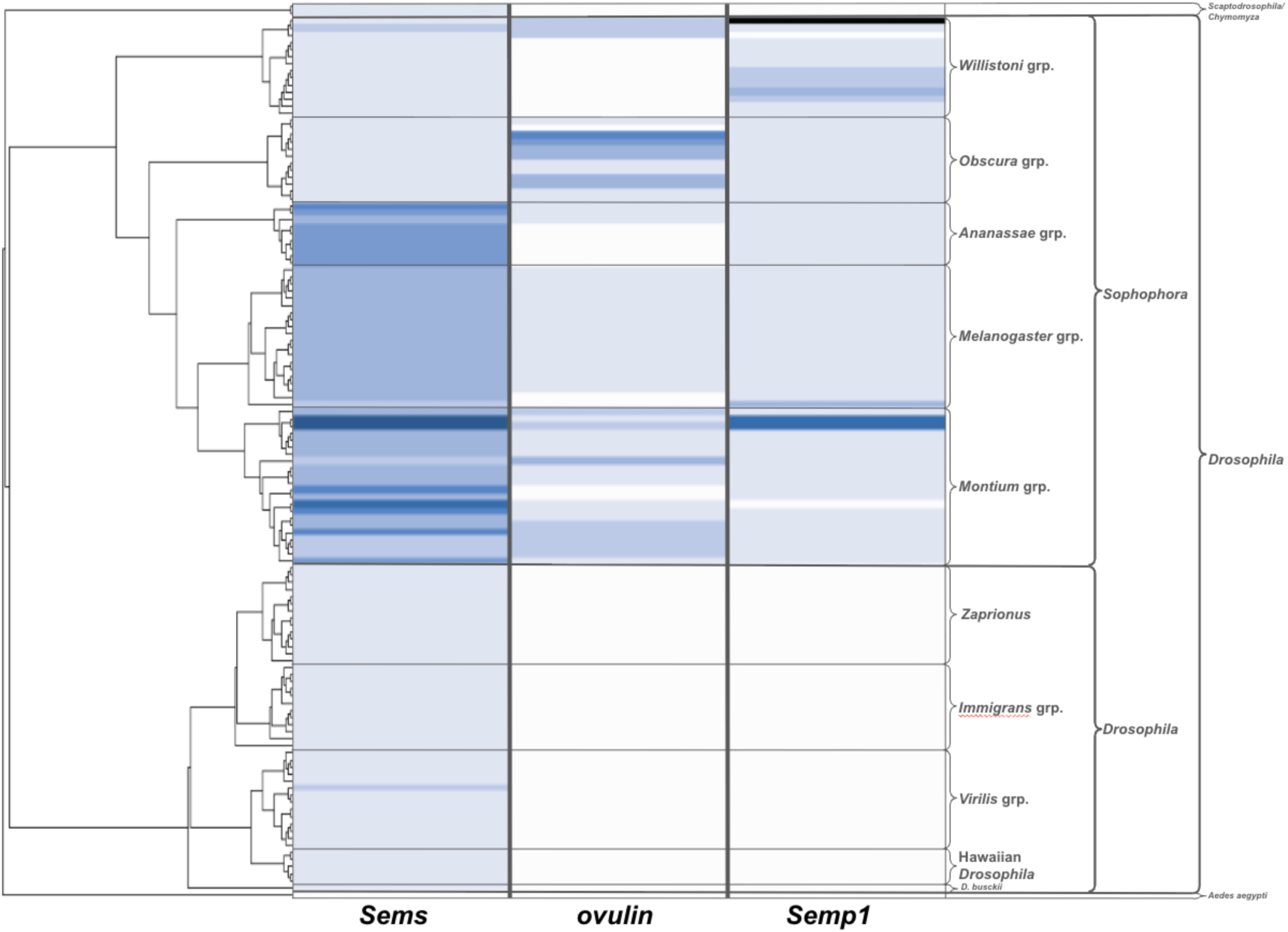
Copy number variation in the genus *Drosophila* of other Sfps functionally related to *Sems*. Copy numbers of *Sems*, *ovulin*, and *Semp1* genes and their close paralogs for 135 Drosophilids inferred via iterative blast and phylogenetic clustering. CNV of *Semp1* is not correlated with *seminase* or *SP*.

**Supplementary Figure 3.**
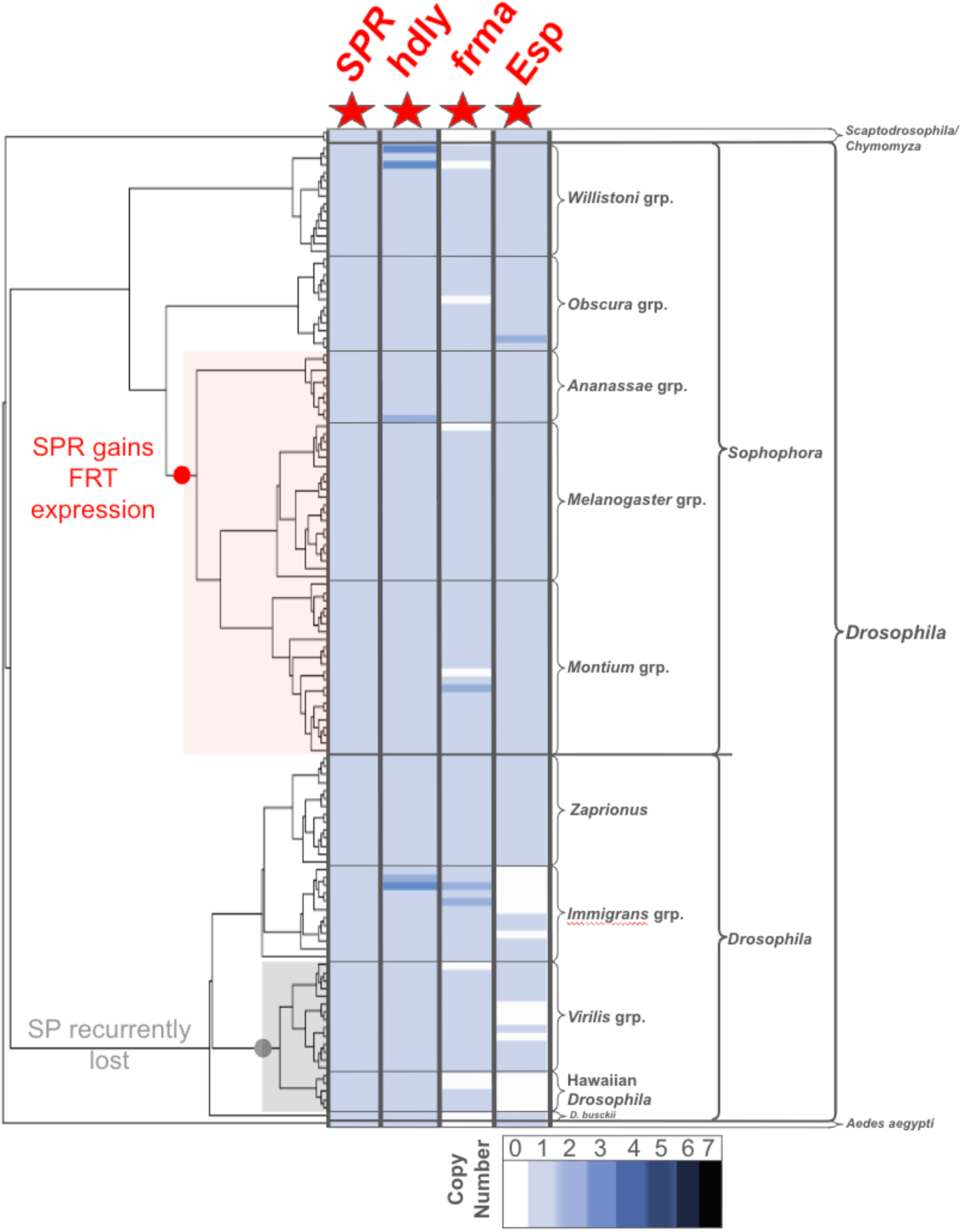
Copy number variation of genes that mediate downstream effects of SP in females and SP’s receptor in the genus *Drosophila*. CNV of SPR from Hopkins *et al* 2024^45^. *Frma*, *Hdly*, and *Esp* CNV determined via same pipeline used on SPN genes. No significant CNV was observed for any of these genes across the genus, although *Hdly* and *Esp* were occasionally lost in some lineages. Therefore, as was the case for SPR, there is no CNV correlation between these genes with each other, *SP*, or the SPN.

