## Supplementary File 2 for "Correlated Gene Copy Number Changes in a Seminal Fluid Protein Network in *Drosophila*"

###############GENERAL NOTES BEFORE STARTING
#Except where noted, all code is Bash (remainder is in R)
#Relevant files are in accompanying folder
#The only parameters that are changed depending on what gene family the pipeline is run on (used in the EMBOSS getORF and bedtools slop command): minimum ORF length (decided on a case-by-case basis; usually ~75% of query legnth), whether to require a start codon (yes for serine proteases, no for all other gene families since they are multi-exon genes and only one exon is used), and whether to collect flanking DNA sequence surrounding hits (bedtools slop).


###############DOWNLOADING GENOMES FROM NCBI
mkdir genomes
cd genomes
###download genomes used in Hopkins et al. 2024 (obtained list of accessions from: https://osf.io/z3a8k). Manually reformatted file and removed entries without accessions and named resultant file "Hopkins_genomes_w_accessions"
export PATH=/programs/ncbi_datasets:$PATH
dos2unix ./Hopkins_genomes_w_accessions
#Create empty files named after each accession to be downloaded 
cat Hopkins_genomes_w_accessions | parallel -j 1 ">{}" &
#iterate over empty files, and download corresponding accession from NCBI datasets
for f in *1*
do
	datasets download genome accession $f --filename "${f}.zip"
done;
#remove empty files
rm *1
rm *2
rm *4
#unzip folders
for f in *.zip
do
	unzip $f -d "${f}_unzipped"
done;
#rename unzipped folder to remove ".zip_unzipped" from folder names
ls -d *unzipped | cut -f 1,2 -d . | parallel -j 10 "mv {}.zip_unzipped {}"
#iterate over folders and genome assembly fasta files into main parent directory
for f in *1
do
	cp ./"${f}"/ncbi_dataset/data/"${f}"/*.fna .
done;
for f in *2
do
	cp ./"${f}"/ncbi_dataset/data/"${f}"/*.fna .
done;


###############BEFORE STARTING: Make directory for analysis and copy all genomes downloaded above to this folder
mkdir <gene_family_name>
cd <gene_family_name>
cp ../genomes/*.fna .
#add genome name to beginning of fasta headers (this is necessary for being able to track which sequences are from which genome throughout the course of the pipeline)
for f in *.fna
do 
	sed -i "s/^>/>${f}_/" "$f"
done;
#remove '.fna' from genome name added to headers above
for f in *.fna
do 
	sed -i "s/.fna//g" "$f"
done;


###############FOR SINGLE EXON GENES ONLY (just used for serine proteases): Blasting to transcriptome to expand initial query; gets the full protein sequence of top 5 blast hits for each query in every Drosophila species with an NCBI predicted transcriptome, then uses these sequences as the query for the main pipeline.
#start in main folder for the gene family (created above)
mkdir making_query_from_transcriptome
cd ./making_query_from_transcriptome
#get all Drosophila NCBI transcriptomes (example below)
cp /fs/cbsuclarkfs1/storage/Drosophila_Genomes1/Drosophila_transcriptomes/*/*.fna .
cp /fs/cbsuclarkfs1/storage/Drosophila_Genomes1/Drosophila_transcriptomes/*/*/*.fna .
#make a query fasta file containing all D. melanogaster protein sequences representing the gene family of interest. Name file 01_Dmel_tblastn_query_seqsfromflybase.
#tblastn Dmel protein seqs for each query to each transcriptome
nohup bash -c 'for f in ./*.fna
do
	tblastn -query 01_Dmel_tblastn_query_seqsfromflybase -subject $f -outfmt 6 -evalue 1e-5 -max_target_seqs 10 -out "${f}_tblastn"
done' &
#Split tblastn output by query, sort by e_val, keep only top 5 hits, combine top 5 hits for all queries
for f in ./*tblastn
do
	awk '{OFS="\t"; print >$1 "_split_by_query"}' $f
	for d in ./*split_by_query
	do
		sort -k 11,11 -g $d >"${d}_sorted"
		head -n 5 "${d}_sorted" >"${d}_sorted_subset"
	done;
	rm *_split_by_query
	rm *sorted
	cat *_sorted_subset >"${f}_top5"
	rm *subset
done;
#combine blast output tables for each genome into one file
cat *_tblastn_top5 > combined_tblastn_table
#get second column of blast output table (will have accession of each hit, so resultant file will be a list of accessions)
awk '{OFS="\t"; print $2}' combined_tblastn_table >accession_list
#remove duplicate accessions from list
awk '!seen[$0]++' accession_list >accession_list_no_dups.txt
#use above file (accession_list_no_dups.txt) to get corresponding protein sequences from batch entrez in fasta format (https://www.ncbi.nlm.nih.gov/sites/batchentrez); name resultant fasta file "all_transcriptome_hits_NCBI_AA.fasta"
#combine with original query to create new query for blasting to all genomes
cat all_transcriptome_hits_NCBI_AA.fasta 01_Dmel_tblastn_query_seqsfromflybase > all_transcriptomes_query.fasta
#copy combined query to parent folder with all genomes
cp all_transcriptomes_query.fasta ../
cp 01* ../
#return to main folder with all genome files and all transcriptome hits
cd ../


###############MAIN PIPELINE
#tblastn protein sequences of all transcriptome hits to each genome (IF USING SINGLE EXON OF MULTI-EXON GENE, SKIP TRANSCRIPTOME STEP AND USE 01_Dmel_tblastn_query_seqsfromflybase AS QUERY IN PLACE OF all_transcriptomes_query.fasta)
nohup sh -c 'ls *.fna | parallel -j 16 "tblastn -query all_transcriptomes_query.fasta -subject {} -outfmt 6 -evalue 1e-5 -max_target_seqs 10 -out {}_tblastn"' &
#Split tblastn output by query, sort by e_val, keep only top 5 hits, combine top 5 hits for all queries
for f in ./*tblastn
do
	awk '{OFS="\t"; print >$1 "_split_by_query"}' $f
	for d in ./*split_by_query
	do
		sort -k 11,11 -g $d >"${d}_sorted"
		head -n 5 "${d}_sorted" >"${d}_sorted_subset"
	done;
	rm *_split_by_query
	rm *sorted
	cat *_sorted_subset >"${f}_top5"
	rm *subset
done; 
#convert subsetted tblastn hits to minimal bed file (i.e. columns are: contig/scaffold; subject start; subject end)
for f in ./*top5
do
	awk '{OFS="\t"; print $2, $9, $10}' $f >"${f}.bed"
done;
#fix cases where end coordinate is lower than start (i.e. reverse hits) so that bedtools can be used
for f in ./*top5.bed
do
	awk '{OFS="\t"; if ($3-$2 > 0) {print}}' $f >"${f}_FORWARD.bed"
	awk '{OFS="\t"; if ($2-$3 > 0) {print}}' $f >"${f}_REVERSE.bed"
	awk ' { OFS="\t"; t = $3; $3 = $2; $2 = t; print; } ' "${f}_REVERSE.bed" >"${f}_REVERSE.bed.rev"
	cat "${f}_FORWARD.bed" "${f}_REVERSE.bed.rev" >"${f}_fixed"
	rm "${f}_FORWARD.bed"
	rm "${f}_REVERSE.bed"
	rm "${f}_REVERSE.bed.rev"
done;
#sort bed file
export PATH=/programs/bedtools2-2.29.2/bin:$PATH
for f in ./*.bed_fixed
do
	sortBed -i $f >"${f}_sorted"
done;
#remove duplicate sequences with bedtools merge (combines all overlapping/bookended hits, which is the default behavior of the function)
for f in ./*.bed_fixed_sorted
do
	bedtools merge -i $f >"${f}_dups_removed"
done;
#make fasta indexes
pip install pyfaidx
for f in ./*.fna
do
	faidx $f -i chromsizes >"${f}_sizes"
done;
#For single exon genes get 200bp on either side of each hit (i.e. set <size_flanking> to 200 in the command below) to ensure entire open reading frame will be recovered. For multi-exon gene where query is single exon, set -b to 0; this will mean the function does nothing to the file but rename it
for f in ./*.fna
do
	bedtools slop -i "${f}_tblastn_top5.bed_fixed_sorted_dups_removed" -g "${f}_sizes" -b <size_flanking> >"${f}_gene_ranges"
done;
#get fasta for genomic sequence corresponding to each hit
for f in ./*.fna
do
	bedtools getfasta -fo "${f}_gene_ranges.fasta" -fi $f -bed "${f}_gene_ranges"
done;
#replace ':' with '_' to fix errors w/ downstream functions
sed -i 's/:/_/g' *gene_ranges.fasta
#get ORFs longer than <minimum_ORF_length> bp; for single exon genes, require ATG start codon at beginning of ORF (-find set to 1); for multiexon genes, do not use this requirement (-find set to 0)
for f in ./*gene_ranges.fasta
do
	singularity run -B $PWD --pwd $PWD /programs/EMBOSS/emboss.sif getorf $f -minsize <minimum_ORF_length> -find <require_start_codon?> "${f}_ORFs"
done;
#if applicable (i.e. there are paralogous outgroups to the gene family of interest), make file containing ~10 closest paralogs and name outgroup_<gene_family_name>
#combine all ORFs for hits across all species as well as the initial query and a file containing outgroups. NOTE: for multi-exon genes, replace "all_transcriptomes_query.fasta" with "01_Dmel_tblastn_query_seqsfromflybase"
cat *ORFs all_transcriptomes_query.fasta outgroup_<gene_family_name> >combined_protein.fasta
#align resultant file with MAFFT using default parameters
export PATH=/programs/mafft/bin:$PATH
mafft --thread 10 combined_protein.fasta > combined_protein_aligned.fasta
#make tree from MAFFT alignment
export PATH=/programs/FastTree-2.1.11:$PATH
nohup sh -c 'FastTree < combined_protein_aligned.fasta > combined_protein.tree' &
#Manually color taxa in figtree by desired orthogroup; note that there may be spurious ORFs in the alignment (especially if not requiring a start codon when using getORF tool). Search for in-group and outgroup sequences on tree to help figure out correct clades to annotate. Save resultant colored tree as combined_protein.nex
#run TREE2FASTA to separate out clades. Before starting, make folder containing the following files; combined_protein.fasta, combined_protein.nex, and TREE2FASTA_v1_6.pl (https://github.com/tomsauv/TREE2FASTA)
perl TREE2FASTA_v1_6.pl combined_protein_R1.nex combined_protein.fasta
#name Tree2Fasta output files <gene_name>_R1 manually (e.g. Sems_R1); if any miscellaneous (i.e. no Dmel sequences included)clades cluster w/ clade of interest but no outgroups, tblastn each ORF in the clade to Dmel transcriptome to confirm the best hit for each member of the miscellaneous clade is within the gene family of interest
#remove any sequences from e.g. transcriptomes/Dmel query (the command keeps only lines that start with >GC, which is only those that came from the genomes as the sequence names in the tree start with the NCBI genome accession of origin)
for f in ./*R1.txt
do 
	grep -e '>GC' $f >"${f}1"
	rm $f
	mv "${f}1" $f
done;
#turn back into bed file (the sequence names in the tree are basically a bed file with "_" as a separator, so this code basically turns each sequence name back into a bed file with an extra column (1st column) containing the name of the genome of origin for each sequence; column 2 is contig/scaffold, column 3 is hit start, and column 4 is hit end).
for f in ./*R1.txt
do 
	rev "$f" | sed 's/_/\t/' | rev >"${f}1"
	rm $f
	mv "${f}1" $f
done;
for f in ./*R1.txt
do 
	rev "$f" | sed 's/-/\t/' | rev >"${f}1"
	rm $f
	mv "${f}1" $f
done;
for f in ./*R1.txt
do 
	rev "$f" | sed 's/_/\t/' | rev >"${f}1"
	rm $f
	mv "${f}1" $f
done;
for f in ./*R1.txt
do 
	rev "$f" | sed 's/_/\t/' | rev >"${f}1"
	rm $f
	mv "${f}1" $f
done;
for f in ./*R1.txt
do 
	rev "$f" | sed 's/_/\t/' | rev >"${f}1"
	rm $f
	mv "${f}1" $f
done;
for f in ./*R1.txt
do
	sed -i "s/>//g" $f
	awk '{OFS="\t"; print $1, $2 "_" $3, $4, $5}' $f >"${f}_fixed"
done;
#Split hits by genome; run this command for each annotated orthogroup (e.g. just change the gene/orthogroup name). Make sure naming matches the names you gave the output files from TREE2FASTA 
awk '{OFS="\t"; print >$1 "_<gene_name>_hitsR1"}' <gene_name>_R1.txt_fixed
#combine all hits to all genes for each genome
ls *.fna | cut -f 1 -d . | parallel -j 10 'cat {.}*hitsR1 >{.}_combined_hitsR1' &
#concatenate columns 1 and 2 (genome name and scaffold/contig name) to turn into normal bed file. Column 1 is contig/scaffold, column 2 is hit start, column 3 is hit end.
for f in ./*_combined_hitsR1
do
	awk '{OFS="\t"; print $1 "_" $2, $3, $4}' $f >"${f}.bed"
done;
#get fasta sequences corresponding to hits in resultant bed files; this bed contains all hits in clade for each species and will be used as a blast query for blasting back to all genomes
export PATH=/programs/bedtools2-2.29.2/bin:$PATH
ls *.fna | cut -f 1 -d . | parallel -j 10 'bedtools getfasta -fo {.}_query -fi {.}*.fna -bed {.}*R1.bed' &
#replace ':' with '_' to fix errors in downstream functions
sed -i 's/:/_/g' *query
#get ORFs longer than <minimum_ORF_length> bp; for single exon genes, require ATG start codon at beginning of ORF (-find set to 1); for multiexon genes, do not use this requirement (-find set to 0). Keep parameters same as last time.
for f in ./*query
do
	singularity run -B $PWD --pwd $PWD /programs/EMBOSS/emboss.sif getorf $f -minsize <minimum_ORF_length> -find <require_start_codon?> "${f}_ORFs"
done;
#combine ORFs of all hits to all genomes as well as starting query into one query file to be used for second round of iterative blast.  NOTE: for multi-exon genes, replace "all_transcriptomes_query.fasta" with "01_Dmel_tblastn_query_seqsfromflybase"
cat *query_ORFs all_transcriptomes_query.fasta > all_hits_query.fasta
#blast all hits from all genomes that cluster with gene family of interest back to all genomes
nohup bash -c "ls *.fna | cut -f 1 -d . | parallel -j 12 'tblastn -query all_hits_query.fasta -subject {.}*fna -outfmt 6 -evalue 1e-5 -max_target_seqs 10 -out {.}_tblastn2'" &
#Split tblastn output by query, sort by e_val, keep only top 5 hits, combine subsetted hits
for f in ./*tblastn2
do
	awk '{OFS="\t"; print >$1 "_split_by_query"}' $f
	for d in ./*split_by_query
	do
		sort -k 11,11 -g $d >"${d}_sorted"
		head -n 5 "${d}_sorted" >"${d}_sorted_subset"
	done;
	rm *_split_by_query
	rm *sorted
	cat *_sorted_subset >"${f}_top5"
	rm *subset
done; 
#convert subsetted tblastn hits to minimal bed file again
for f in ./*tblastn2_top5
do
	awk '{OFS="\t"; print $2, $9, $10}' $f >"${f}.bed"
done;
#fix cases where end coordinate is lower than start (i.e. reverse hits) 
for f in ./*tblastn2_top5.bed
do
	awk '{OFS="\t"; if ($3-$2 > 0) {print}}' $f >"${f}_FORWARD.bed"
	awk '{OFS="\t"; if ($2-$3 > 0) {print}}' $f >"${f}_REVERSE.bed"
	awk ' { OFS="\t"; t = $3; $3 = $2; $2 = t; print; } ' "${f}_REVERSE.bed" >"${f}_REVERSE.bed.rev"
	cat "${f}_FORWARD.bed" "${f}_REVERSE.bed.rev" >"${f}_fixed"
	rm "${f}_FORWARD.bed"
	rm "${f}_REVERSE.bed"
	rm "${f}_REVERSE.bed.rev"
done;
#combine new hits with first round of hits
ls *.fna | cut -f 1 -d . | parallel -j 10 'cat {.}*tblastn2_top5.bed_fixed {.}*dups_removed >{.}_R1R2' &
#sort bed file
export PATH=/programs/bedtools2-2.29.2/bin:$PATH
for f in ./*R1R2
do
	sortBed -i $f >"${f}_sorted"
done;
#remove duplicates with bedtools merge (default is to combine all overlapping/bookended hits)
for f in ./*R1R2_sorted
do
	bedtools merge -i $f >"${f}_dups_removed"
done;
#For single exon genes get 200bp on either side of each hit (i.e. set <size_flanking> to 200 in the command below) to ensure entire open reading frame will be recovered. For multi-exon gene where query is single exon, set -b to 0; this will mean the function does nothing to the file but rename it
ls *.fna | cut -f 1 -d . | parallel -j 10 'bedtools slop -i {.}*R1R2_sorted_dups_removed -g {.}*sizes -b <size_flanking> >{.}_R2_gene_ranges' &
#get fasta
ls *.fna | cut -f 1 -d . | parallel -j 10 'bedtools getfasta -fo {.}_R2_gene_ranges.fasta -fi {.}*.fna -bed {.}*R2_gene_ranges' &
#replace ':' with '_' to fix errors w/ downstream functions
sed -i 's/:/_/g' *R2_gene_ranges.fasta
#get ORFs longer than <minimum_ORF_length> bp; for single exon genes, require ATG start codon at beginning of ORF (-find set to 1); for multiexon genes, do not use this requirement (-find set to 0). Keep parameters same as last time.
for f in ./*R2_gene_ranges.fasta
do
	singularity run -B $PWD --pwd $PWD /programs/EMBOSS/emboss.sif getorf $f -minsize <minimum_ORF_length> -find <require_start_codon?> "${f}_ORFs"
done;
###NOTE: from this point forward, the rest of pipeline can be run several different ways (different combinations of aligners (MAFFT/PROMALS3D) and tree-building programs (fasttree, raxml)). When I do this, I make the alignment and tree using one set of methods, complete the rest of the pipeline, then transfer all files created after this point to a folder named after the aligner/tree program used (e.g. R2_MAFFT_fasttree) and start again from this point with the next method.
----------------------------------------------------------------------------------------------------------------------V1 (MAFFT + fasttree)
#combine all ORFs across all spp.
cat *R2_gene_ranges.fasta_ORFs outgroup* 01* >combined_protein_R2.fasta
#WARNING: CHECK THAT CONACATENATION DIDN'T ADD ONE OF THE FASTA HEADERS TO THE END OF ANOTHER ENTRY INSTEAD OF IN ITS OWN LINE
#align resultant file 
export PATH=/programs/mafft/bin:$PATH
mafft --thread 10 combined_protein_R2.fasta > combined_protein_R2_aligned.fasta
#make tree from alignment
export PATH=/programs/FastTree-2.1.11:$PATH
nohup bash -c 'FastTree < combined_protein_R2_aligned.fasta > combined_protein_R2.tree' &
#Manually color taxa in figtree by desired orthogroup; note that there may be spurious ORFs in the alignment (especially if not requiring a start codon when using getORF tool). Search for in-group and outgroup sequences on tree to help figure out correct clades to annotate. Save resultant colored tree as combined_protein.nex
perl TREE2FASTA_v1_6.pl combined_protein_R2.nex combined_protein_R2.fasta
#name Tree2Fasta output files {gene}_R2 manually (e.g. Sems_R2)
----------------------------------------------------------------------------------------------------------------------V2 (PROMALS3D + RAxML-ng)
mkdir R2_PROMALS3D_raxml
cd R2_PROMALS3D_raxml
cp ../combined_protein_R2.fasta .
##align with PROMALS3D
#PROMALS3D only accepts names up to 25 characters, so need to replace each name with a number in the input fasta, then replace the numbers with the correct names in the alignment output
sed 's/ //g' combined_protein_R2.fasta >combined_protein_R2.fasta_PROMALS3D
sed -i 's/\[.*//' combined_protein_R2.fasta_PROMALS3D
grep -e ">" combined_protein_R2.fasta_PROMALS3D >name_key_col1
sed -i 's/>//g' name_key_col1
awk -i ord -v OFS=, '
    function randint(n) { return int(n*rand()) }
    BEGIN { srand(); A = ord("A"); Z = ord("Z") }
    NR == 1 { printf("%s,CASENUMBER\n", $0); next }
    { printf "%s,%c%c%.5d\n", $0, A + randint(Z-A+1), A + randint(Z-A+1), randint(100000) }' name_key_col1 >name_key
while read -r line;do
    #echo "${line#*,}"
	#echo "${line%,*}"
	sed -i "s/${line%,*}/${line#*,}/g" combined_protein_R2.fasta_PROMALS3D
done < name_key
#now use combined_protein_R2.fasta_PROMALS3D file as input on PROMALS3D aligner webserver (http://prodata.swmed.edu/promals3d/promals3d.php). Name output combined_protein_filtered.fasta_PROMALS3D_output and put back in same folder
while read -r line;do
    #echo "${line#*,}"
	#echo "${line%,*}"
	sed -i "s/${line#*,}/${line%,*}/g" combined_protein_R2.fasta_PROMALS3D_output_processed
done < name_key
###Making tree with raxml-ng
export PATH=/programs/raxml-ng_v1.2.0:$PATH
#calculate appropriate number of cores for tree/bootstrapping
raxml-ng --parse --msa combined_protein_R2.fasta_PROMALS3D_output_processed --model LG --prefix T1
#make tree
nohup sh -c 'raxml-ng --msa combined_protein_R2.fasta_PROMALS3D_output_processed --model LG --prefix T3 --threads <number_cores> --seed 6' &
#calculate bootstraps
nohup sh -c 'raxml-ng --bootstrap --msa combined_protein_R2.fasta_PROMALS3D_output_processed --model LG --prefix T7 --seed 569 --threads <number_cores>' &
#put bootstrap values on tree
raxml-ng --support --tree T3.raxml.bestTree --bs-trees T7.raxml.bootstraps --prefix T14 --threads 1 --bs-metric tbe
#Manually color taxa in figtree by desired orthogroup; note that there may be spurious ORFs in the alignment (especially if not requiring a start codon when using getORF tool). Search for in-group and outgroup sequences on tree to help figure out correct clades to annotate. Save resultant colored tree as combined_protein.nex
perl TREE2FASTA_v1_6.pl combined_protein_R2.nex combined_protein_R2.fasta_PROMALS3D_output_processed
#name Tree2Fasta output files {gene}_R2 manually (e.g. Sems_R2)
----------------------------------------------------------------------------------------------------------------------V3 (MAFFT + RAxML-ng)
mkdir R2_MAFFT_raxml
cd R2_MAFFT_raxml
cp ../*R2*/combined_protein_R2_aligned.fasta .
cp ../combined_protein_R2_aligned.fasta .
###Making tree with raxml-ng
export PATH=/programs/raxml-ng_v1.2.0:$PATH
#calculate appropriate number of cores for tree/bootstrapping
raxml-ng --parse --msa combined_protein_R2_aligned.fasta --model LG --prefix T1
#make tree
nohup sh -c 'raxml-ng --msa combined_protein_R2_aligned.fasta --model LG --prefix T3 --threads <number_cores> --seed 6' &
#calculate bootstraps
nohup sh -c 'raxml-ng --bootstrap --msa combined_protein_R2_aligned.fasta --model LG --prefix T7 --seed 569 --threads <number_cores>' &
#put bootstrap values on tree
raxml-ng --support --tree T3.raxml.bestTree --bs-trees T7.raxml.bootstraps --prefix T14 --threads 1 --bs-metric tbe
----------------------------------------------------------------------------------------------------------------------V4 (PROMALS3D + iqTree)
mkdir R2_PROMALS3D_iqTree
cd R2_PROMALS3D_iqTree
cp ../R2*/combined_protein_R2.fasta_PROMALS3D_output_processed .
nohup sh -c '/programs/iqtree-2.2.2.6-Linux/bin/iqtree2 -s combined_protein_R2.fasta_PROMALS3D_output_processed -B 1000 -T AUTO' &
----------------------------------------------------------------------------------------------------------------------V5 (MAFFT + iqTree)
mkdir R2_MAFFT_iqtree
cd R2_MAFFT_iqtree
cp ../*R2*/combined_protein_R2_aligned.fasta .
cp ../combined_protein_R2_aligned.fasta .
nohup sh -c '/programs/iqtree-2.2.2.6-Linux/bin/iqtree2 -s combined_protein_R2_aligned.fasta -B 1000 -T AUTO' &
----------------------------------------------------------------------------------------------------------------------V6 (MAFFT + FastTree)
mkdir R2_PROMALS3D_FastTree
cd R2_PROMALS3D_FastTree
cp ../R2*/combined_protein_R2.fasta_PROMALS3D_output_processed .
#make tree from alignment
export PATH=/programs/FastTree-2.1.11:$PATH
nohup bash -c 'FastTree < combined_protein_R2.fasta_PROMALS3D_output_processed > combined_protein_R2.fasta_PROMALS3D_output_processed.tree' &
----------------------------------------------------------------------------------------------------------------------CONTINUE FROM HERE 
#remove any sequences from query (i.e. keep only hits from genome)
for f in ./*R2.txt
do 
	grep -e '>GC' $f >"${f}1"
	rm $f
	mv "${f}1" $f
done;
#turn back into bed file (with extra column (column 1) with genome name)
for f in ./*R2.txt
do 
	rev "$f" | sed 's/_/\t/' | rev >"${f}1"
	rm $f
	mv "${f}1" $f
done;
for f in ./*R2.txt
do 
	rev "$f" | sed 's/-/\t/' | rev >"${f}1"
	rm $f
	mv "${f}1" $f
done;
for f in ./*R2.txt
do 
	rev "$f" | sed 's/_/\t/' | rev >"${f}1"
	rm $f
	mv "${f}1" $f
done;
for f in ./*R2.txt
do 
	rev "$f" | sed 's/_/\t/' | rev >"${f}1"
	rm $f
	mv "${f}1" $f
done;
for f in ./*R2.txt
do 
	rev "$f" | sed 's/_/\t/' | rev >"${f}1"
	rm $f
	mv "${f}1" $f
done;
for f in ./*R2.txt
do
	sed -i "s/>//g" $f
	awk '{OFS="\t"; print $1, $2 "_" $3, $4, $5}' $f >"${f}_fixed"
done;
#Split hits by genome; run this command for each annotated orthogroup (e.g. just change the gene/orthogroup name). Make sure naming matches the names you gave the output files from TREE2FASTA 
awk '{OFS="\t"; print >$1 "_<gene_name>_hitsR2"}' <gene_name>_R2.txt_fixed
#remove carryover from query
rm Dmel*
rm Dele*
#concatenate genome name and contig/scaffold name columns to make bed file 
for f in ./*_hitsR2
do
	awk '{OFS="\t"; print $1 "_" $2, $3, $4}' $f >"${f}.bed"
done;
#get fasta for each gene in each species
export PATH=/programs/bedtools2-2.29.2/bin:$PATH
ls *<gene_name>_hitsR2.bed | cut -f 1 -d . | parallel -j 2 'bedtools getfasta -fo {.}_query_<gene_name> -fi {.}*.fna -bed {.}*<gene_name>_hitsR2.bed' &
#combine fastas for each 
cat *_query_<gene_name> >combined_reciprocal_query_<gene_name>.fasta
#replace ':' with '_' to fix errors with downstream functions
sed -i 's/:/_/g' *reciprocal_query*
#get ORFs
for f in ./*reciprocal_query*
do
	singularity run -B $PWD --pwd $PWD /programs/EMBOSS/emboss.sif getorf $f -minsize <minimum_ORF_length> -find <require_start_codon?> "${f}_ORFs"
done;
#tblastn to Dmel (check reciprocal best blast)
for f in ./*reciprocal_query*_ORFs
do
	tblastn -query $f -subject Dmel_rna.fna -outfmt 6 -evalue 1e-5 -max_target_seqs 1 -out "${f}_results"
done;
#Split tblastn output by query, sort by e_val, keep only best hits, combine subsetted hits
for f in ./*reciprocal_query*_ORFs*results
do
	awk '{OFS="\t"; print >$1 "_split_by_query"}' $f
	for d in ./*split_by_query
	do
		sort -k 11,11 -g $d >"${d}_sorted"
		head -n 1 "${d}_sorted" >"${d}_sorted_subset"
	done;
	rm *_split_by_query
	rm *sorted
	cat *_sorted_subset >"${f}_best_hit"
	rm *subset
done;
#get table containing name of each query and of corresponding best hit
for f in ./*reciprocal_query*_ORFs*results*best_hit
do
	awk '{OFS="\t"; print $1, $2}' $f >"${f}_list"
done;
#get only second column and remove duplicates (just unique best hits in the Dmel transcriptome)
for f in ./*reciprocal_query*_ORFs*results*best_hit_list
do
	awk '{OFS="\t"; print $2}' $f >"${f}_unique"
done;
for f in ./*reciprocal_query*_ORFs*results*best_hit_list_unique
do
	awk '!seen[$0]++' $f >"${f}1"
	mv "${f}1" $f
done;
#check in output that all accessions are expected Dmel best hit to confirm reciprocal best blast
#NOTE: spurious ORFs (i.e. DNA hit is real but got multiple ORFs, some in the wrong frame) may hit a closely related gene; check any weird cases manually and make sure they have another ORF from the same DNA sequence that hits the right gene.
#NOTE: tandem dups (e.g. intr/CG34295) may have both genes/ORFs in DNA sequence since I used bedtools merge to get rid of duplicates by combining overlapping or bookended sequences. This happens because I just got ORFs separately for each <gene_name>_hitsR2.bed file and e.g. intr_hitsR2.bed may share a region with CG34295_hitsR2.bed (i.e. because the ORFs of both genes were in a single DNA hit). DOES NOT AFFECT COUNTS OR CAUSE ANY OTHER ISSUES OUTSIDE OF RECIPROCAL BLASTING.
#combine all hits to all genes for each genome
ls *.fna | cut -f 1 -d . | parallel -j 10 'cat {.}*hitsR2 >{.}_combined_hitsR2' &
#combine genome and contig/scaffold columns to make bed file
for f in ./*_combined_hitsR2
do
	awk '{OFS="\t"; print $1 "_" $2, $3, $4}' $f >"${f}.bed"
done;
#Make table of copy number counts for each orthogroup in each genome
wc -l *hitsR2 >countsR2.txt
#WARNING SHORTEN GENE NAMES TO NOT BE SPEAD ACROSS MULTIPLE UNDERSCORES (e.g. replace "misc1_Ser12_paralog" w/ "misc1")
###replace genome names in counts file with species name
#transfer file titled "species_name_genome_key" to analysis folder
#replace genome accessions in counts table with corresponding genome name based on values in key file
dos2unix  species_name_genome_key
awk '{OFS="\t"; print $2, $1}' species_name_genome_key >species_name_genome_key1
sed -i 's/_/\t/2' countsR2.txt 
sed -i 's/\.1//g' species_name_genome_key1 
sed -i 's/\.2//g' species_name_genome_key1 
sed -i 's/\.1\t/\t/g' countsR2.txt 
sed -i 's/\.2\t/\t/g' countsR2.txt
awk '{OFS="\t"; print $2, $3, $1}' countsR2.txt >countsR2.txt1
sed -i 's/_combined/	combined/g' countsR2.txt1
awk 'FNR==NR{a[$1]=$2;next} {$0=a[$1]" "$0 }1' species_name_genome_key1 countsR2.txt1 >countsR2_renamed.txt
awk '{OFS="\t"; print $4, $1, $2, $3}' countsR2_renamed.txt >countsR2_renamed_final.txt
#make sure to remove extra (e.g. title) rows at top and bottom of file
#make data matrix for visualization (species by gene)
sed -i 's/combined/combined_/g' countsR2_renamed_final.txt
rev countsR2_renamed_final.txt | sed 's/_/\t/' | rev >countsR2_renamed_final.txt1
rm countsR2_renamed_final.txt
mv countsR2_renamed_final.txt1 countsR2_renamed_final.txt
rev countsR2_renamed_final.txt | sed 's/_/\t/' | rev >countsR2_renamed_final.txt1
rm countsR2_renamed_final.txt
mv countsR2_renamed_final.txt1 countsR2_renamed_final.txt
sed -i 's/combined\t/placeholder\tcombined/g' countsR2_renamed_final.txt
awk '{OFS="\t"; print $2, $5, $1}' countsR2_renamed_final.txt >final_counts_table
#manually delete duplicate species (Drosophila_paulistorum, Drosophila_immigrans, Drosophila_willistoni, Zaprionus_indianus x2, Drosophila_teissieri)
#also delete any cases where accession fails to be replaced by genome name (if applicable; in my experience, only happens with D. paulistorum)
###RUN REMAINDER OF PIPELINE IN R
#import "final_counts_table" as <gene_family_name>_final_counts
library(tidyr)
library(stringr)
library(dplyr)
<gene_family_name>_count_matrix <- pivot_wider(data = <gene_family_name>_final_counts, V1, names_from = V2, values_from = V3)
<gene_family_name>_count_matrix <- <gene_family_name>_count_matrix %>% mutate_all( ~replace(., lengths(.)==0, 0))
<gene_family_name>_count_matrix[is.na(<gene_family_name>_count_matrix)] <- 0
rownames(<gene_family_name>_count_matrix) <- <gene_family_name>_count_matrix$V1
#manually clean up data matrix (e.g. rename species to match tree etc.); name
#make heatmap in R with rows ordered such that they match the species on the tree
library(ape)
library(TreeTools)
library(RColorBrewer)
setwd("C:/PATH/TO/FOLDER/WITH/PHYLOGENY/FILE")
Drosophila_tree = read.tree(file = "Species_phylogeny.txt")
<gene_family_name>_count_matrix <- <gene_family_name>_count_matrix
<gene_family_name>_count_matrix$rownames <- rownames(<gene_family_name>_count_matrix)
<gene_family_name>_count_matrix <- <gene_family_name>_count_matrix[match(Drosophila_tree$tip.label, <gene_family_name>_count_matrix$rownames),]
<gene_family_name>_count_matrix$rownames <- NULL
heatmap(as.matrix(<gene_family_name>_count_matrix[]), Rowv = NA, Colv = NA, scale = "none", col = hcl.colors(50, "Oslo"))


###############SPECIES TREE-GENE TREE RECONCILIATION WITH GENERAX
sed -i 's/>//g' <gene_family_name>_R2.txt
#Get sequences for gene of interest from second (and final) sequence collection tree
export PATH=/programs/seqkit-0.15.0:$PATH
seqkit grep -f <gene_family_name>_R2.txt combined_protein_R2.fasta -o <gene_family_name>_R2.fasta
#rename sequences to pass through PROMALS3D 
sed 's/ //g' <gene_family_name>_R2.fasta ><gene_family_name>_R2.fasta_PROMALS3D
sed -i 's/\[.*//' <gene_family_name>_R2.fasta_PROMALS3D
grep -e ">" <gene_family_name>_R2.fasta_PROMALS3D >name_key_col1
sed -i 's/>//g' name_key_col1
awk -i ord -v OFS=, '
    function randint(n) { return int(n*rand()) }
    BEGIN { srand(); A = ord("A"); Z = ord("Z") }
    NR == 1 { printf("%s,CASENUMBER\n", $0); next }
    { printf "%s,%c%c%.5d\n", $0, A + randint(Z-A+1), A + randint(Z-A+1), randint(100000) }' name_key_col1 >name_key
sed -i 's/ //g' name_key
while read -r line;do
    #echo "${line#*,}"
	#echo "${line%,*}"
	sed -i "s/${line%,*}/${line#*,}/g" <gene_family_name>_R2.fasta_PROMALS3D
done < name_key
#remove any duplicate sequences (has to match in both name AND sequence to be removed, so this function doesn't remove duplicates different names/identifiers)
export PATH=/programs/seqkit-0.15.0:$PATH
seqkit rmdup < <gene_family_name>_R2.fasta_PROMALS3D > <gene_family_name>_R2.fasta_PROMALS3D_NoDups
rm <gene_family_name>_R2.fasta_PROMALS3D
mv <gene_family_name>_R2.fasta_PROMALS3D_NoDups <gene_family_name>_R2.fasta_PROMALS3D
#now use combined_protein_filtered.fasta_PROMALS3D file as input on PROMALS3D aligner webserver (http://prodata.swmed.edu/promals3d/promals3d.php). Name output combined_reciprocal_query_<gene_family_name>.fasta_ORFs_PROMALS3D_output and put back in same folder
#import combined_reciprocal_query_<gene_family_name>.fasta_ORFs_PROMALS3D into geneious and visually check alignment. For any outlying sequences, run InterProScan and keep them if they are Serine Proteases and remove them if not (if they are not, they are probably spurious ORFs). Remove all gaps with less than 50% coverage across sequences (also in geneious). Name resultant file combined_reciprocal_query_<gene_family_name>.fasta_ORFs_PROMALS3D_output_processed, then run code below:
while read -r line;do
    #echo "${line#*,}"
	#echo "${line%,*}"
	sed -i "s/${line#*,}/${line%,*}/g" <gene_family_name>_R2.fasta_PROMALS3D_output_processed
done < name_key
#Remove duplicate sequences (by sequence this time regardless of name); save duplicates in output file titled "duplicated.fa"
export PATH=/programs/seqkit-0.15.0:$PATH
seqkit rmdup -s -d duplicated.fa < <gene_family_name>_R2.fasta_PROMALS3D_output_processed > <gene_family_name>_R2.fasta_PROMALS3D_output_processed_dups_removed
#The goal of this step is to find and remove any technical duplicates due to assembly errors (e.g. duplicated contig containing gene of interest). Go through "duplicated.fa" file manually and check that all duplicate sequences are from different genomes (i.e. same amino acid sequence in different species)--as long as this is the case, then none of the duplicates are spurious and you can add them back into the alignment. For any duplicates from the same species, check the DNA sequence of the gene and flanking regions on both contigs/scaffolds to determine whether the identical copies are likely technical or biological and retain/discard the sequences accordingly.
#Replace genome accessions with species names
sed -i 's/_/./g' <gene_family_name>_R2.fasta_PROMALS3D_output_processed_dups_removed 
cp ../*/species_name_genome_key_reconciliation_new .
dos2unix species_name_genome_key_reconciliation_new
while read -r line;do
    #echo "${line#*,}"
	#echo "${line%,*}"
	sed -i "s/${line#*,}/${line%,*}/g" <gene_family_name>_R2.fasta_PROMALS3D_output_processed_dups_removed 
done < species_name_genome_key_reconciliation_new
#Remove any sequences that were not renamed (this would mean the genome was not in the "species_genome_key_reconciliation" key file, meaning that either A) it wasn't in Ben Hopkins Sex Peptide protein fasta file or B) multiple genomes representing the same species were present in the analysis and thus the duplicates can be ignored)
#RENAME D. melanogaster sequences to match naming conventions
sed -i 's/Dmel/D.MELANOGASTER_/g' <gene_family_name>_R2.fasta_PROMALS3D_output_processed_dups_removed 
#Build the input gene tree for GeneRax (set <substitution_model> to either LG or LG+G, depending on which model will be used downstream in GeneRax; we performed sepearate RAxML-ng/GeneRax runs for each gene family with each of these two substitution models).
export PATH=/programs/raxml-ng_v1.2.0:$PATH
raxml-ng --parse --msa <gene_family_name>_R2.fasta_PROMALS3D_output_processed_dups_removed --msa-format fasta --model LG+G --prefix T1
nohup sh -c 'raxml-ng --msa <gene_family_name>_R2.fasta_PROMALS3D_output_processed_dups_removed --model LG+G --prefix T3 --threads 4 --seed 6' &
#Species tree-gene tree reconciliation with GeneRax. See discussion of the substitution model in the RAxML-ng comment directly above. 
cp ../*/Kim_* .
export PATH=/programs/GeneRax-2.1.3/bin:$PATH
nohup sh -c 'generax -f families_<gene_family_name> -s Kim_2024_subset.tree -r UndatedDL' &


################Creation of bed files for manually checking synteny. Each file contains the best hit in each genome for 10 genes flanking each locus of interest.
#input is fasta of protein sequence for 10 genes flanking (5 on each side) gene of interest; file titled e.g. "Protease_Synteny_Genes.txt"
#tblastn to all genomes
nohup sh -c 'ls *.fna | parallel -j 12 "tblastn -query <gene_name>_Synteny_Genes.txt -subject {} -outfmt 6 -evalue 1e-5 -max_target_seqs 10 -out {}_synt_tblastn"' &
#get only top hit for each query
for f in ./*tblastn
do
	awk '{OFS="\t"; print >$1 "_split_by_query"}' $f
	for d in ./*split_by_query
	do
		sort -k 11,11 -g $d >"${d}_sorted"
		head -n 1 "${d}_sorted" >"${d}_sorted_subset"
	done;
	rm *_split_by_query
	rm *sorted
	cat *_sorted_subset >"${f}_top_hit"
	rm *subset
done; 
#convert subsetted tblastn hits to minimal bed (fourth column is feature name)
for f in ./*tblastn_top_hit
do
	awk '{OFS="\t"; print $2, $9, $10, $1}' $f >"${f}.bed"
done;

#fix cases where end coordinate is lower than start (i.e. reverse hits)
for f in ./*tblastn_top_hit.bed
do
	awk '{OFS="\t"; if ($3-$2 > 0) {print}}' $f >"${f}_FORWARD.bed"
	awk '{OFS="\t"; if ($2-$3 > 0) {print}}' $f >"${f}_REVERSE.bed"
	awk ' { OFS="\t"; t = $3; $3 = $2; $2 = t; print; } ' "${f}_REVERSE.bed" >"${f}_REVERSE.bed.rev"
	cat "${f}_FORWARD.bed" "${f}_REVERSE.bed.rev" >"${f}_fixed"
	rm "${f}_FORWARD.bed"
	rm "${f}_REVERSE.bed"
	rm "${f}_REVERSE.bed.rev"
done;

#split bed file by gene for which flanking genes were blasted
for f in ./*tblastn_top_hit.bed_fixed
do
	grep -e '<gene_name>' $f >"${f}_<gene_name>"
done;
#the resultant bed file can then be used to manually check synteny and confirm loss events.
