## Supplementary File 3 for "Correlated Gene Copy Number Changes in a Seminal Fluid Protein Network in *Drosophila*"

### Supplemental methods and guide to data files:

**SECTION 1: Annotation of orthologous and paralogous sequences in 135 Drosophilds to assess copy number variation.**

**NOTE:** All relevant files for each run of the pipeline are found within folders named after the gene family (e.g. Serine_Proteases). Code for the pipeline with detailed comments is found in the main folder in a file called “SPN_paper_pipeline_final.txt”. While most of the pipeline does not vary between runs, a few parameters differ between gene families to accommodate differences in gene structure. The parameters for each run are found in Supplemental File 1, Table 3. Information about the generation and naming of additional relevant files will be laid out below.

1. **For single exon genes** (only used for serine protease clade): Expand initial query by blasting gene family of interest to all *Drosophila* NCBI predicted transcriptomes
    a. Starting with the amino acid sequence of all *Drosophila melanogaster* genes nested within the gene family of interest, we used tblastn (e-value cutoff: 1e-5) to identify the five best hits by e-value for each query in the 35 *Drosophila* species for which NCBI predicted transcriptomes were available. Duplicate accessions were removed from the hit list and the full protein sequences were obtained from NCBI via Batch Entrez. These sequences were then combined with the initial *D. melanogaster* protein sequence query and used as the input for the main annotation pipeline.
    b. Relevant files:
        i. 01_Dmel_tblastn_query_seqsfromflybase: initial protein query in fasta format
        ii. all_transcriptomes_query.fasta: the expanded query used as the input for step 2

2. First round of iterative blast and phylogenetic clustering.
    a. Starting with either A) for single-exon genes, the expanded query created in the previous step or B) for multi-exon genes, the amino acid sequence of the longest exon containing a conserved functional domain for each *Drosophila melanogaster* gene nested within the gene family of interest[1], we used tblastn (e-value cutoff: 1e-5) to identify the five best hits (by e-value) for each query in 135 Drosophilid genomes. Using bedtools, we then merged overlapping hits and obtained the corresponding genomic DNA sequences[2]. Open reading frames were predicted using the EMBOSS getorf tool[3]. The resultant ORFs were

---

[1]Care was also taken to ensure that the intron-exon structure of the gene region chosen as query was conserved across all members of the gene family of interest as well as in the closest paralogous outgroups in *D. melanogaster*.

[2] For single-exon genes, since we require an ATG start codon for predicted ORFs, 200 base pairs flanking each hit were also obtained to ensure recovery of the start codon. For multi-exon genes, whether or not flanking regions were collected varied by gene family (see Supplemental File 1, Table 3)

[3] Minimum ORF length was set to ~75% of the ORF length of the shortest query sequence; an ATG start codon was required for single-exon gene families but not for multi-exon families.

combined with the initial query file, the expanded query file (single-exon genes only), as well as a file containing the protein sequences of the ~10 closest paralogs to the gene family of interest. This combined fasta was then aligned using MAFFT and a protein phylogeny inferred using FastTree. Clades of orthologous proteins were then manually annotated on the tree using FigTree; the corresponding sequence names were obtained from the tree using TREE2FASTA.

b. Relevant files:
    i. 01_Dmel_Seqs_From_flybase: initial protein query in fasta format
    ii. all_transcriptomes_query.fasta: the expanded query used as the input for step 2 (NOTE: only for single-exon gene families)
    iii. outgroups_<gene_family>: a FASTA file containing the amino acid sequence of the ~10 closest *D. melanogaster* paralogs to the gene family of interest (for all gene families except hdly, since it is an orphan gene).
    iv. combined_protein_R1.fasta: the ORFs of all hits hits from the first round of blasting combined with the initial query and outgroup sequences (these sequences are used to build the first tree)
    v. combined_protein_aligned_R1.fasta: the above file aligned using MAFFT
    vi. combined_protein_R1.tree: a phylogeny built from the above amino acid alignment using FastTree
    vii. combined_protein_R1.nex: the above tree, with orthogroups manually annotated (by taxon color) in FigTree
    viii. <gene_name>_R1.txt: text files containing a list of all sequences belonging to each orthogroup manually annotated on the tree in the file above (sequence names match preceding alignments/trees)

c. Relevant folders:
    i. bed_fasta_R1: folder containing the following files
        1. <genome_accession>_combined_hitsR1.bed: bed files for hits that cluster with the gene family of interest for each genome
        2. <genome_accession>_query: FASTA file containing DNA sequences of all hits
        3. <genome_accession>_query_ORFs: FASTA file containing amino acid sequences of all hits

3. Second round of iterative blast and phylogenetic clustering.
    a. The approach is identical to Step 2 above except that the tblastn query is all hits that convincingly cluster with the gene family of interest from the first round of iterative blasting; the tools and settings used were otherwise the same, except where noted. Briefly, the top 5 best hits (by e-value) for each query were collected from 135 genomes and combined with the first round of hits, overlapping hits were combined, corresponding DNA sequences were obtained, and ORFs were predicted. All ORFs, queries, and outgroup sequences were combined and protein trees were inferred using six different combinations of aligners and phylogenetic inference tools: MAFFT + FastTree; MAFFT +

RAxML-ng; MAFFT + IQ-TREE 2; PROMALS3D + FastTree; PROMALS3D + RAxML-ng; PROMALS3D + IQ-TREE 2. Clades of orthologous proteins were again manually annotated on the MAFFT/FastTree tree using FigTree and the corresponding sequence lists obtained using TREE2FASTA[4]. Comparison of tree topology and robustness of annotated clades across methods was examined and is shown in supplemental figures 1-6.

b. Relevant files:
    i. all_hits_query.fasta: the query for the second tblastn search. Contains the protein sequences of the genes of interest from *D. melanogaster* as well as all hits from the first round of iterative blasting to the genomes that cluster convincingly with the gene family of interest.
    ii. combined_protein_R2.fasta: the ORFs of all hits hits from the first and second rounds of blasting combined with the initial query and outgroup sequences (these sequences are used to build the first tree)
    iii. combined_protein_R2_aligned.fasta: the above FASTA file aligned using MAFFT
    iv. combined_protein_R2.fasta_PROMALS3D_output_processed: the above FASTA file aligned with PROMALS3D
    v. combined_protein_R2_MAFFT_FastTree: a phylogeny built from the above MAFFT amino acid alignment using FastTree
    vi. combined_protein_R2_MAFFT_RAxML: a phylogeny built from the above MAFFT amino acid alignment using RAxML-ng
    vii. combined_protein_R2_MAFFT_IQ-TREE: a phylogeny built from the above MAFFT amino acid alignment using IQ-TREE 2
    viii. combined_protein_R2_PROMALS3D_RAxML: a phylogeny built from the above PROMALS3D amino acid alignment using RAxML-ng
    ix. combined_protein_R2_PROMALS3D_IQ-TREE: a phylogeny built from the above PROMALS3D amino acid alignment using IQ-TREE 2
    x. combined_protein_R2_<aligner>_<tree_program>.nex: one of the above trees with orthogroups annotated (by taxon color) in FigTree. The orthogroups shown were initially annotated on the MAFFT/FastTree tree then mapped onto the other trees to allow easy visual comparison.
    xi. <gene_name>_R2.txt: text files containing a list of all sequences comprising each orthogroup (sequence names match preceding alignments/trees).

c. Relevant folder:
    i. bed_fasta_R2
        1. <genome_accession>_<gene_name>_hitsR2.bed: bed files for each genome separated by gene

[4]For *hdly* and *frma*, to deal with a small number of spurious ORFs incorrectly nested within clade of interest on tree with long branch lengths, additional manual filtering was performed on the sequence list to remove hits that both had little/no similarity to the *D. melanogaster hdly* or *frma* transcripts (as assessed via tblastn to the Dmel transcriptome) and were derived from the same DNA sequence as a genuine hit but in the wrong frame (4 sequences removed for *hdly* and 6 for *frma*). Additionally, for *frma*, a single hit that formed an outgroup to all of the others was removed as it did not pass reciprocal best blast.

2. <genome_accession>_query_<gene_name>: a FASTA file containing the DNA sequences of all hits from the second and final round of sequence collection separated by both orthogroup and genome
3. combined_reciprocal_query_<gene_name>.fasta: a FASTA file containing the DNA sequences of all hits from the second and final round of sequence collection separated by orthogroup
4. combined_reciprocal_query_<gene_name>.fasta_ORFs: a FASTA file containing the amino acid sequences of all hits from the second and final round of sequence collection separated by orthogroup

4. Confirming that the best tblastn hit in the *D. melanogaster* transcriptome for all orthologs/paralogs annotated is the expected gene.
    a. Hits from the second and final round of iterative tblastn sequence collection were separated by orthogroup, the corresponding DNA sequences were obtained, and the ORFs were predicted. We then used tblastn to find the best hit for each query in the NCBI *D. melanogaster* predicted transcriptome. The output was then checked to confirm that all queries (i.e. hits validated by phylogenetic clustering) hit the expected *D. melanogaster* gene[5].
    b. Relevant files:
        i. combined_reciprocal_query_<gene_name>.fasta_ORFS: a FASTA file containing the amino acid sequences of all hits from the second and final round of sequence collection separated by orthogroup. These files were used as the tblastn queries for identifying best hits in the *D. melanogaster* transcriptome.
        ii. combined_reciprocal_query_<gene_name>.fasta_ORFs_results_best_hit_list: a table of the best hits for each member of a given orthogroup. Column 1 is the query sequence (one of the hits annotated by the pipeline) and column 2 is the NCBI accession of the best hit.
        iii. combined_reciprocal_query_<gene_name>.fasta_ORFs_results_best_hit_list_unique: a list of unique values for accession from the above file.
5. Visualization of copy number variation:
    a. A table of counts from step 3 was processed in R to first transform it into a matrix and then visualize the copy number variation as a heatmap. The following packages were used; tidyr, stringr, dplyr, ape, TreeTools, and RColorBrewer
    b. Relevant files:

[5] There were two types of cases where the best hit was not the expected gene: 1) cases where spurious ORFs (i.e. ORFs in the wrong frame) weakly hit a paralogous Dmel gene; 2) cases where two tandem hits (e.g. intr/CG34295) are combined into a single DNA range, so both ORFs are present (note that this issue only arises in the reciprocal best blast step and does not bias copy number estimation). In both cases, in addition to the spurious hit, there should be a second ORF predicted from the same DNA sequence that does hit the expected gene. In our dataset, all cases where an unexpected best hit was recovered were determined to have arisen from one of these processes.

        i. countsR2_renamed_final.txt: A table of copy number counts. Column 1 is count, column 2 is the species name, column 3 is the genome accession, and column 5 is the gene name (ignore columns 4 and 6).
        ii. count_matrix.csv: A matrix of copy number counts made from the above table
        iii. Phylogeny_for_visualization.txt: the phylogeny used to arrange the rows (species) in the CNV heatmap figures

6. Generation of bed files for checking synteny to confirm losses:
    a. We manually created a FASTA file containing the amino acid sequence of ~10 *D. melanogaster* genes flanking each genomic locus containing at least one gene of interest. In selecting flanking genes, we chose longer and functionally characterized genes where possible. We then used tblastn to search for the best hit (e-value) for each of these flanking genes in all the genomes, turned the hit table into a bed file showing the location of each hit, and used these bed files to manually confirm loss events via synteny.
    b. Relevant folders:
        i. Synteny_genes:
            1. Relevant files:
                a. <gene_family>_synteny_genes.fasta: query file containing *D. melanogaster* protein sequences of ten genes flanking each locus of interest
                b. <genome_accession>.fna_synt_tblastn_top_hit.bed_fixed_<genes_of_interest_at_locus>: a bed file containing the location of the best hit in the associated genome for each of the flanking genes in the query file above

Miscellaneous supplemental files for running pipeline:

- species_name_genome_key: tsv table containing correspondence between genome accession numbers and species names
- Hopkins_genomes_w_accessions: list of assembly accession numbers used in this study (originally obtained from Hopkins et al. (2024))

**Technical issues:**

- Spurious ORFs (i.e. ORFs in the wrong frame) sometimes cause issues with alignment/tree inference, especially when using PROMALS3D as the aligner. This problem was most pronounced for *hdly*, likely because it is a rapidly evolving orphan gene and the exon analyzed is short. See supplemental File 4 for a comparison of the performance of different combinations of aligners and tree inference methods for each gene family. Note that as the minimum allowable ORF length is increased, the number of spurious ORFs in the dataset should be reduced, but the rate of false negatives (i.e. inferring a loss where none actually occurred) may go up if changes in gene structure have occurred.
- The estimation of copy number may be biased by structural changes in some paralogs. For example:

- The gain of an intron interrupting the section of the gene used as query in some lineages might result in failure to recover some orthologs/paralogs
- The duplication of a domain/exon within a gene might result in inflated copy number estimates if the entire section of the gene used as query duplicates

- For some gene families, the longest conserved exon is relatively short, which may limit the power to resolve phylogenetic relationships between hits.

**SECTION 2: Gene tree-species tree reconciliation with GeneRax (and subsequent correlation of gains and losses between gene families)**

1. Gene tree-species tree reconciliation:
   a. We used GeneRax to perform gene tree-species tree reconciliation to infer the where on the species tree gene gains and losses occurred for all orthogroups. Briefly, the amino acid sequences for each orthogroup (annotated by the main pipeline) were aligned using PROMALS3D. Sites with less than 50% occupancy were then masked using Geneious. The alignment was subsequently manually checked for spurious ORFs (i.e. secondary ORFs in the wrong frame)[6] and technical duplicates (though no such duplicates were found)[7]. Any offending sequences were removed from the alignment. The starting gene tree was then inferred from the resultant alignment using RAxML-ng and the starting species tree was taken from Hopkins et al. (2024) and subsetted to remove species not present in our analysis (leaving 131 spp.); these files alongside the manually filtered PROMALS3D protein alignment were used as the inputs for GeneRax. In all cases, only duplications and losses (i.e. no transfers) were allowed (--rec-model set to UndatedDL). Correlations between rates of events between gene families were computed in R.
   b. Relevant folders:
      1. <orthogroup>_<substitution_model>_GeneRax
         a. Files:
            i. Drosophila_phylogeny_subset.txt: the species tree from Hopkins et al. (2024) subsetted such that only species considered in the analysis presented here are included
            ii. combined_reciprocal_query_<gene_name>.fasta_ORFs_PROMALS3D_output_processed_dups_removed: masked (>50% occupancy) PROMALS3D

---

[6] The alignment was checked visually for outlying sequences. We removed any outliers that satisfied both of the following two conditions: 1) when blasted (tblastn) to the *D. melanogaster* transcriptome, no significant similarity was found; and 2) there was a second non-outlying ORF recovered from the same DNA sequence.

[7] All cases where two sequences in the alignment had exactly the same amino acid sequence (identified by seqkit rmdup) were manually checked to ensure that either 1) they were from two different species and thus were not spurious or 2) if they were from the same species, that the corresponding DNA sequence of the identical hits and their flanking regions were differentiated enough that they are unlikely to reflect technical duplication due to misassembly.

alignment of all orthogroup sequences after manually filtering

iii. T3.raxml.bestTree: RAxML-ng tree made from the above alignment
iv. families_<orthogroup>: configuration file for GeneRax run specifying input files
v. <orthogroup>_reconciliation.xml: the reconciliated gene tree superimposed on the species tree. Can be visualized using RecPhyloXML
vi. <orthogroup>_speciesEventCounts.txt: The unprocessed table produced by the GeneRax run containing the inferred counts for duplications and losses
vii. gains_losses_table_for_correlation.csv: A manually restructured and cleaned-up table of event counts for all gene families; the input file used for assessing the correlations

**SECTION 3:** Miscellaneous analyses

1. Confirming that concerted loss events shown in Figure 2 are not the product of assembly errors by blasting lost genes to the associated raw reads (only doing for short read assemblies of dubious quality; for long read-based assemblies, syntenic confirmation of loss is sufficient)
   a. For each concerted loss, start with the full length amino acid sequences of all SPN genes in *D. melanogaster*
   b. tblastn each protein sequence to the raw reads associated with the species affected by each putative loss event as well as several close outgroup species (of equal phylogenetic distance from query) that retain the genes
   c. Compare the distribution of e-values obtained between genomes that have and haven't lost the genes
      i. The logic of looking at the distribution rather than e.g. the e-value of just the top hit is that if the loss stems from a pseudogenization event, there may still be some short reads that hit well, but there should be many fewer and the e-values should decay *much* faster moving down the hit list than for an intact gene
      ii. The distribution of e-values obtained will obviously be affected by the length of reads, depth of sequencing, length of query, and level of conservation of the gene of interest, so all of these things must be considered in interpreting the results. Fortunately, most of the short read data used for the Montium group is from a single sequencing project that was internally consistent methodologically, which simplifies things considerably.

iii. *Seminase*, *frma*, and *hdly* can be used (roughly) as controls for depth of sequencing etc. since they are present in almost every species and thus should provide some idea of what to expect from an intact gene in that particular sequencing run
