## Supplementary File 4 for "Correlated Gene Copy Number Changes in a Seminal Fluid Protein Network in *Drosophila*"

### Slide 1
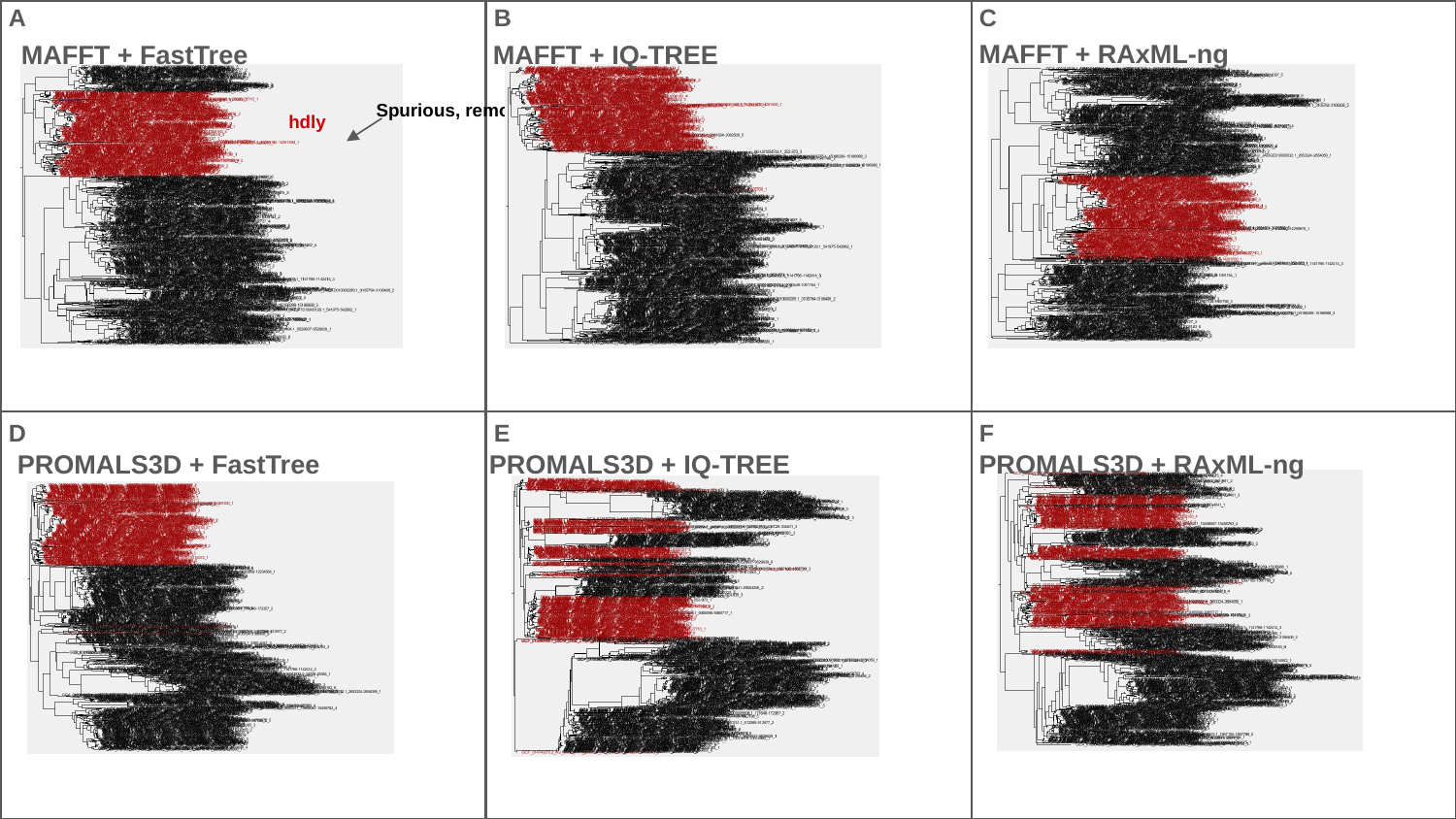

A
B
C
MAFFT + RAxML-ng
MAFFT + FastTree
MAFFT + IQ-TREE
Spurious, removed
hdly
D
E
F
PROMALS3D + FastTree
PROMALS3D + IQ-TREE
PROMALS3D + RAxML-ng

### Slide 2
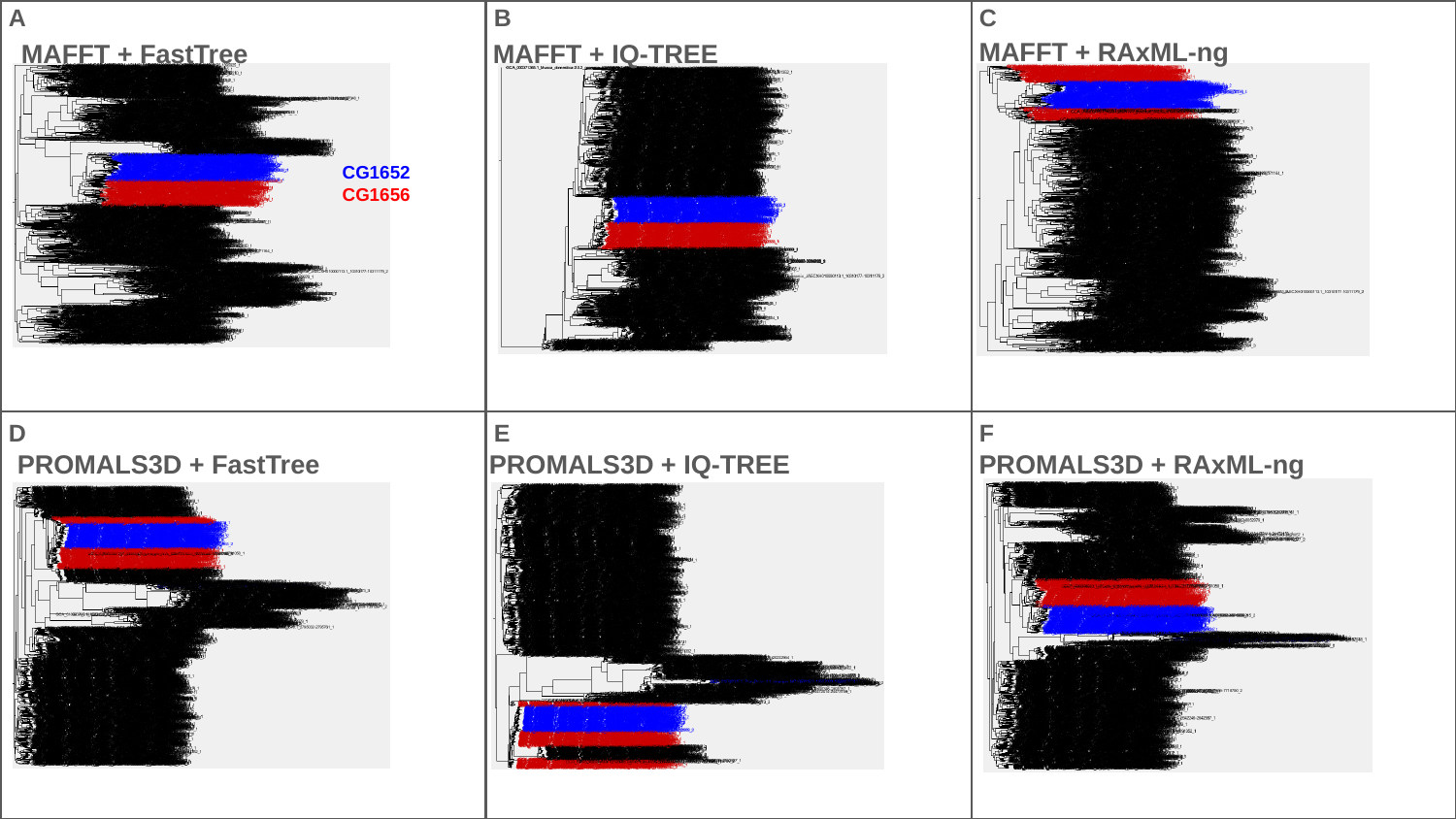

A
B
C
MAFFT + RAxML-ng
MAFFT + FastTree
MAFFT + IQ-TREE
CG1652
CG1656
D
E
F
PROMALS3D + FastTree
PROMALS3D + IQ-TREE
PROMALS3D + RAxML-ng

### Slide 3
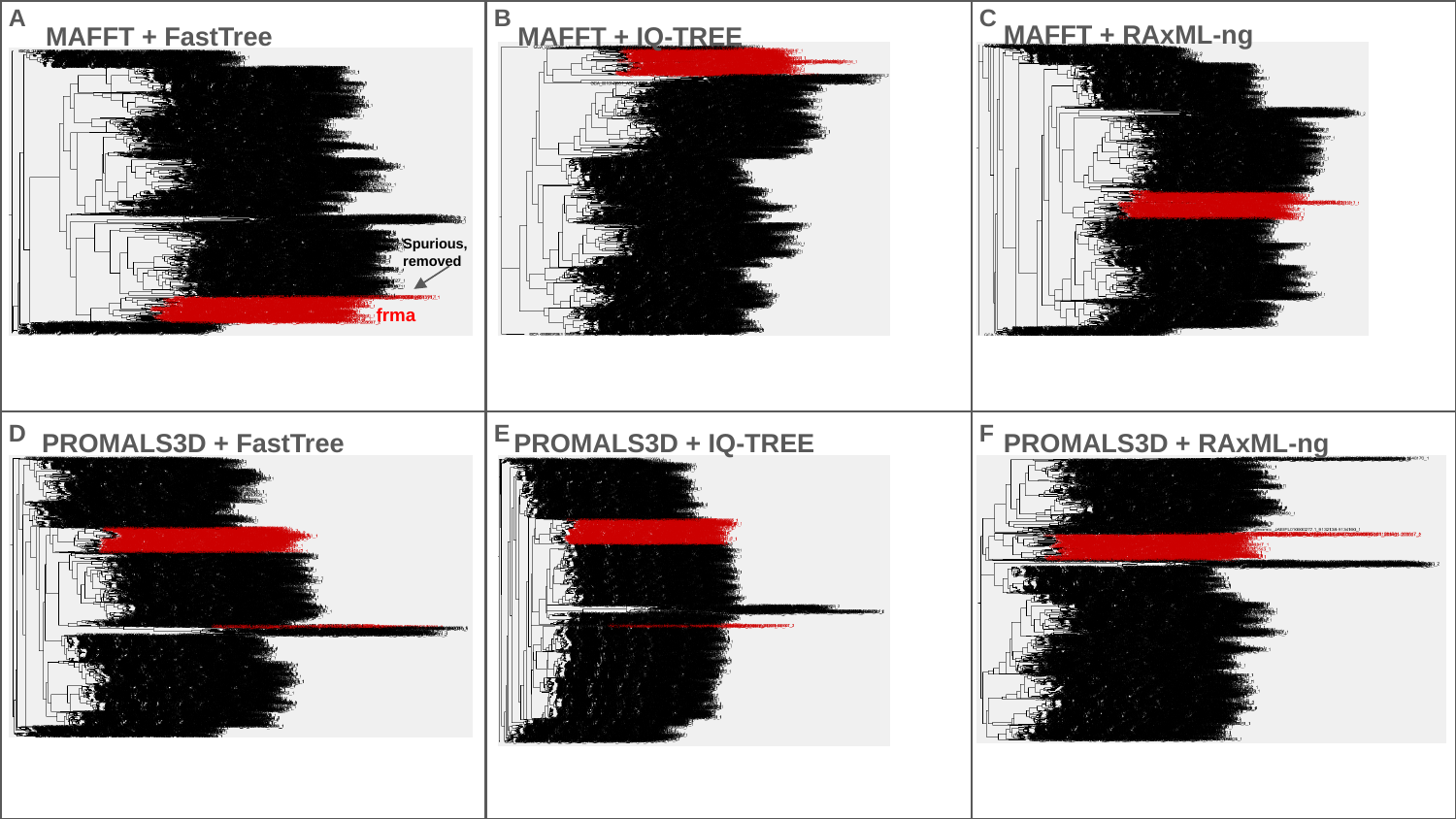

A
B
C
MAFFT + RAxML-ng
MAFFT + FastTree
MAFFT + IQ-TREE
Spurious, removed
frma
D
E
F
PROMALS3D + FastTree
PROMALS3D + IQ-TREE
PROMALS3D + RAxML-ng

### Slide 4
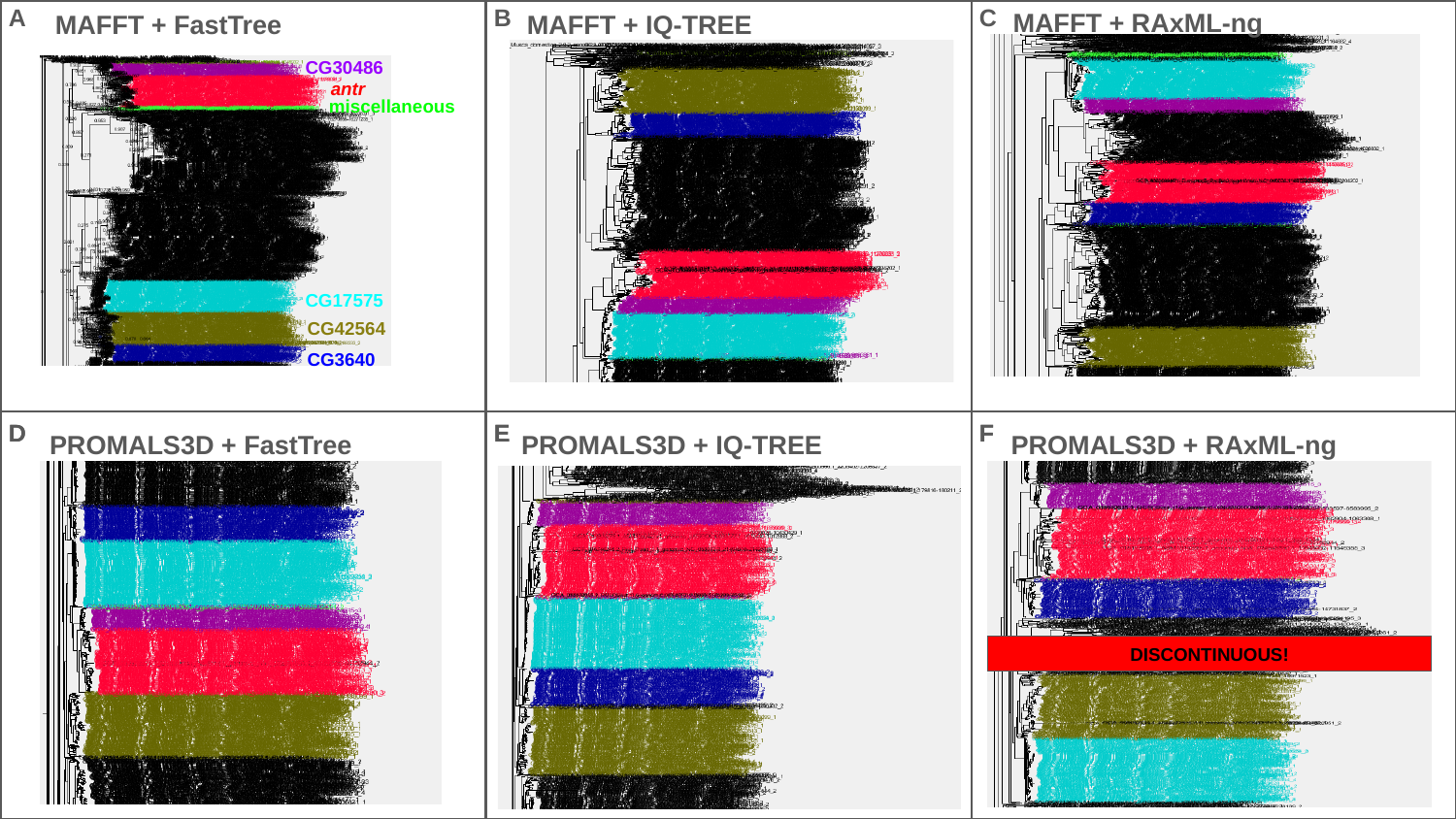

A
B
C
MAFFT + RAxML-ng
MAFFT + FastTree
MAFFT + IQ-TREE
CG30486
 antr
miscellaneous
CG17575
CG42564
CG3640
D
D
E
E
F
F
PROMALS3D + FastTree
PROMALS3D + IQ-TREE
PROMALS3D + RAxML-ng
DISCONTINUOUS!

### Slide 5
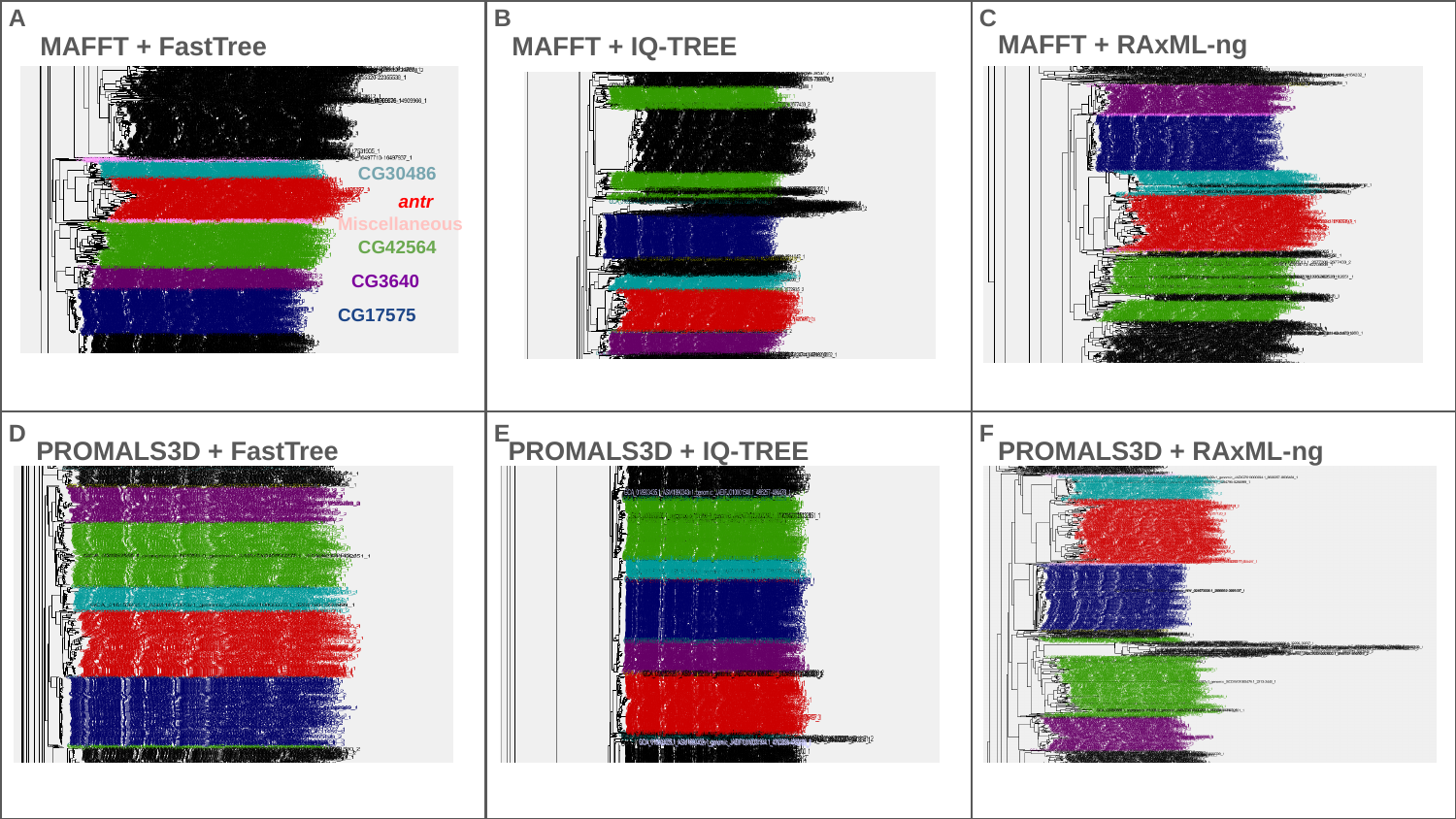

A
B
C
MAFFT + RAxML-ng
MAFFT + FastTree
MAFFT + IQ-TREE
CG30486
antr
Miscellaneous
CG42564
CG3640
CG17575
D
E
F
PROMALS3D + FastTree
PROMALS3D + IQ-TREE
PROMALS3D + RAxML-ng

### Slide 6
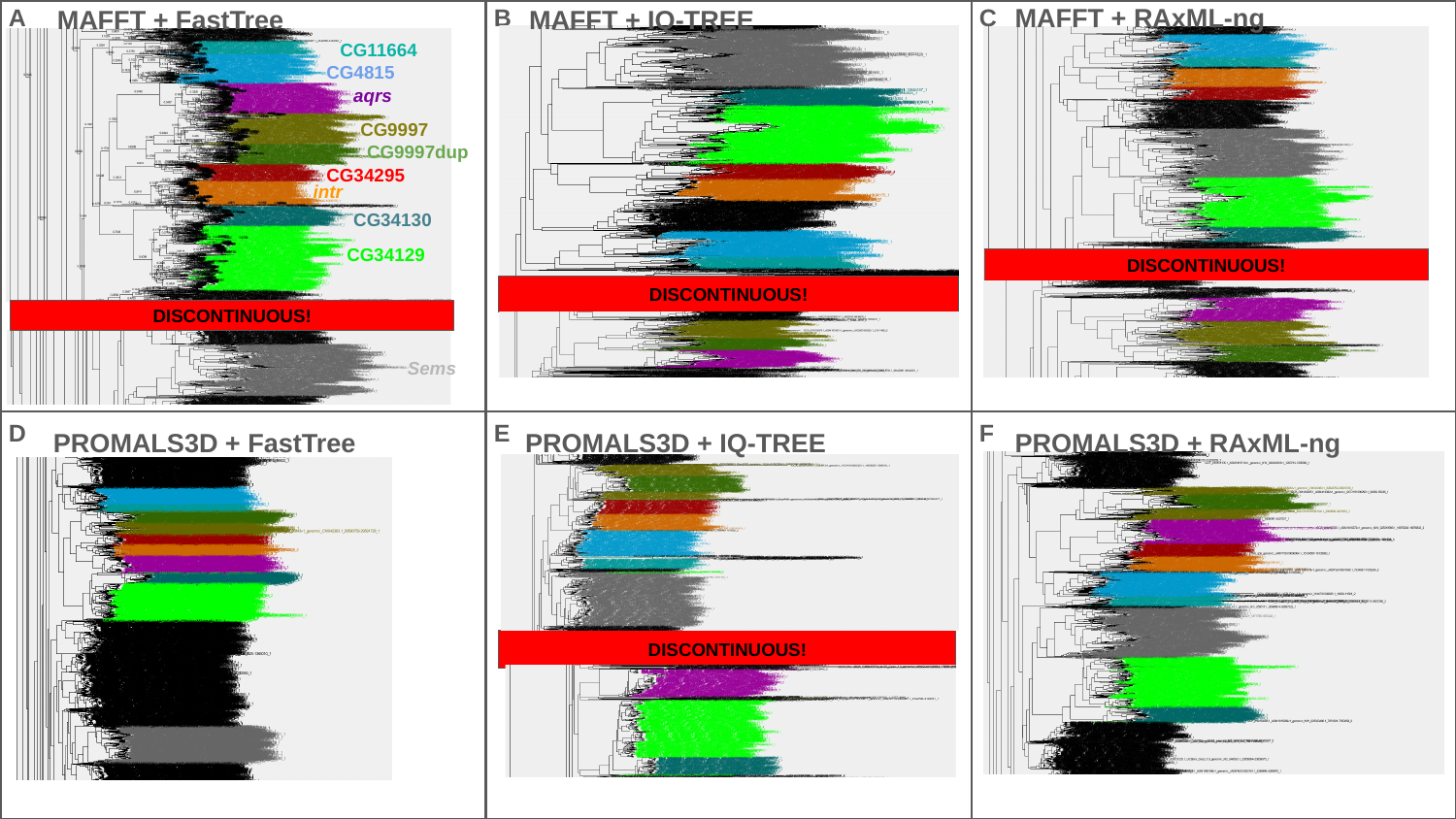

MAFFT + RAxML-ng
A
B
C
MAFFT + FastTree
MAFFT + IQ-TREE
CG11664
CG4815
aqrs
CG9997
CG9997dup
CG34295
intr
CG34130
CG34129
DISCONTINUOUS!
DISCONTINUOUS!
DISCONTINUOUS!
Sems
D
E
F
PROMALS3D + FastTree
PROMALS3D + IQ-TREE
PROMALS3D + RAxML-ng
DISCONTINUOUS!
