## Supplementary File 5 for "Correlated Gene Copy Number Changes in a Seminal Fluid Protein Network in *Drosophila*"

Table 1, Fly stocks used

| **Short name, label** | **Genotype** | **Identifier/RRID** |
|---|---|---|
| *RNAi line 1 against CG42564* | *y,sc,v,sev; P{TRiP.HMC06173}attP40* | BDSC_65910 |
| *RNAi line 2 against CG42564* | *w-;;P{GD6109}v14158* | SCR_013805: VDRC Id 14158 |
| *tub-GAL4* | *y,w; P{tubP-GAL4}LL7* | BDSC_5138 |
| *UAS-Dcr2* | *w1118, P{UAS-dicer2, w[+]}* | SCR_013805: VDRC Id 60007 |
| Wild type CS | Canton S wild type strain | Laboratory of B. Dickson |

Table 2, Genotypes of experimental flies

| **Figure label** | **Full genotype** |
|---|---|
| *tub-GAL4 x RNAi BDSC_65910* | *+; P{TRiP.HMC06173}attP40/+; tub-GAL4/+* |
| *RNAi BDSC_65910* control | *+; P{TRiP.HMC06173}attP40/+; +/+* |
| *tub-GAL4 x RNAi VDRC_14158* | *w-,UAS-Dcr2; +/+; tub-GAL4/P{GD6109}v14158* |
| *RNAi VDRC_14158* control | *+; +/+; P{GD6109}v14158/+* |
| *tub-GAL4* control | *w-,UAS-Dcr2; +/+; tub-GAL4/+* |
